# Analysis of human colostrum reveals differential co-occurrence networks of metabolites, microbiota and cytokines in maternal obesity

**DOI:** 10.1101/2025.03.27.645789

**Authors:** July S. Gámez-Valdez, Karina Corona-Cervantes, Erick S. Sanchez-Salguero, Mario R. Alcorta-García, Claudia N. Lopez-Villaseñor, Rommel A. Carballo-Castañeda, Aldo Moreno-Ulloa, Victor J. Lara-Diaz, Marion E. G. Brunck, Cuauhtémoc Licona-Cassani

**Affiliations:** Centro de Biotecnología FEMSA, Escuela de Ingeniería y Ciencias, Tecnologico de Monterrey, N.L. México; Unidad de Biología Integrativa, Institute for Obesity Research, Tecnologico de Monterrey, N.L. México; Unidad de Biología Experimental, Institute for Obesity Research, Tecnologico de Monterrey, N.L. México; Hospital Regional Materno Infantil, Servicios de Salud de Nuevo León, OPD, Ciudad Guadalupe, N.L. México; Hospital San José TECSALUD, Monterrey, N.L. México; Escuela de Medicina y Ciencias de la Salud, Tecnológico de Monterrey, N.L. México; Departamento de Innovación Biomédica, Centro de Investigación Científica y de Educación Superior de Ensenada, Baja California (CICESE), Ensenada, México; Faculty of Medicine, The University of New South Wales, Sydney, Australia

## Abstract

Breastmilk is essential for neonatal development, particularly in seeding the gut microbiota and modulating the maturing immune system. This proof-of-concept study explores the systemic nature of colostrum and the influence of maternal obesity on co-occurrences of colostrum bioactives. Using 16S-rRNA sequencing, untargeted metabolomics, and cytokines quantification, we analyzed co-occurring elements in the colostrum of mothers with normal weight (18.5 < BMI < 25) or obesity (BMI > 30). We identified 5 different co-occurrence networks, characterized by positive correlations of taxonomically related bacteria. Our integrative analysis reveals that *Aeromonadaceae*, *Xanthomonadaceae* and *Staphylococcaceae* negatively correlate with proinflammatory cytokines TNF-α, IL-6, and IL-12p70 in the colostrum of mothers with obesity (WO). Additionally, lipid mediators, including 15-HEDE and LysoPC(16:00), were associated with cytokines IL-10 and IL-8, and microbiota taxa *Burkholderiaceae*, *Beijerinckiaceae* and *Planococcaceae*–firstly reported in colostrum from mothers WO. Our findings suggest a pervasive regulation of bioactives in the colostrum of mothers WO. This may have implications for a distinctive neonatal intestine development.

## Introduction

Breastmilk is a complex and dynamic fluid that supplies hydration, nutrients, and bioactives essential for the optimal growth and development of neonates.^1^ The bioactive components of breastmilk regulate vital functions, including passive immunity to pathogens, tolerance to the nascent microbiota, and overall processes necessary to mature the intestinal system.^2,3^ Multiple maternal factors have been linked to alterations in breastmilk composition including BMI, age, ethnicity and diet.^4–6^

Most of the current literature draws direct correlations between maternal physiological states and alterations in individual milk components.^7^ While informative, this approach fails to capture a more comprehensive breadth of how maternal condition correlates to breastmilk bioactives. This becomes evident when considering breastmilk as a complex and dynamic ecological system. As an example, within the maternal gut during pregnancy, a clear biological network emerges, wherein bacterial fermentation produces short-chain fatty acids (SCFAs), triggering the proliferation of regulatory T (Treg) cells and modulating the balance between proinflammatory and anti-inflammatory cytokines, thereby promoting an immunotolerant environment that allows the establishment of commensal bacteria.^8^ Understanding breastmilk as a biological system and how it is influenced by maternal health status provides valuable insights into its potential impacts on neonatal health.^9,10^

One in three pregnant women worldwide is affected by obesity. This metabolic disorder has been correlated with alterations at the systemic level, including chronic inflammation. The breastmilk bioactive components of mothers with obesity are also altered.^5,11,12^ For instance, the breastmilk microbiota from mothers with obesity has increased proportions of *Staphylococcus* and *Corynebacterium*.^13,14^ Yet, little is known about possible co-dependances of these alterations with other breastmilk bioactive components in obesity. The breastmilk microbiota is a living component that produces metabolites and other molecules while utilizing maternal factors. Therefore, it is crucial to examine the interactions among soluble bioactives together with the microbiota composition.^15–17^ Approaching breastmilk, and more importantly, colostrum – the initial biological message that lactating mother has formulated for the neonate – as an ecosystem opens avenues to explore the complexity and dynamism of its interacting components.

Using co-occurrence networks to analyze patterns of interactions among bioactives represents a promising approach for exploring the structure of complex biological systems and their interactions with the host.^18^ This approach provides valuable insights on potential interactions that are difficult to detect through characterizing bioactives independently or by using general ecological metrics such as alpha/beta diversity.^19^ Here, we integrate colostrum cytokine concentrations, microbiota composition and metabolites in the context of maternal obesity as an approach to identify distinctive co-occurring networks. By exploring these interconnections, we provide a proof of concept of the systemic nature of colostrum and begin to dimension the complex biological networks at play within this microenvironment.

## Materials and methods

### Experimental model and study participants details

This is an observational study conducted in accordance with the ethical principles outlined in the Declaration of Helsinki. The protocol was approved by the IRB at Escuela de Medicina y Ciencias de la Salud, Tecnologico de Monterrey, with the ID: P000487; it has been registered at Clinical Trials with the ID: NCT04812847. Every participant was provided with details regarding the study, and written consent was acquired from each of them, ensuring the confidentiality of the personal information.

Healthy breastfeeding mothers who had given birth to full-term infants in the Hospital Regional Materno Infantil were recruited between November 2020 and July 2022. The inclusion criteria for participants encompassed mothers between 20 and 32 years, with confirmed residency within the metropolitan area of Monterrey. The exclusion criteria included a history of antibiotic usage in the 3 months prior delivery, a prolonged antibiotic exposure exceeding 3 weeks at any stage of pregnancy, antibiotic requirement for more than 24 hours post-delivery, immunosuppressive, or immunomodulatory corticosteroid therapy, history of feeding disorders or bariatric surgery, experienced delayed lactogenesis or issues with milk supply, pre-term delivery or if required intensive care post-delivery.

Pre-pregnancy BMI was determined by the clinician and reported in the corresponding clinical records. Participants were divided into two groups according to the World Health Organization (WHO) classification of body mass index (BMI)^20^: women with obesity (“WO”; BMI□≥□30 kg/m^2^; n=28) and mothers with normal weight (“NW”; BMI□≤□25 kg/m^2^; n=20).

### Sample collection and processing

Colostrum samples were collected within the first 48 hours post-partum from a total of 48 mothers by a trained medical team, utilizing sterile gloves for the process. Following a gentle cleansing of the areola with sterile water, when possible, 3 mL of colostrum was obtained by manual expression into a sterile 15 mL polypropylene tube. To ensure the purity of the sample, the initial drops were discarded. After collection, the samples were promptly transported to the laboratory and stored at -20 °C until subsequent processing.

### Cytokine measurement

Due to limited sample volume availability, 47 colostrum samples (NW = 20 and WO = 27) were centrifuged (3000 x g, 15 minutes) to remove any cellular structure. Cytokines were quantified in the supernatant using the LEGEND plex® kit (inflammation cytokines markers thirteen, Bio Legend Cat #740808) which includes the IL-1β, TNFα, IL-6, IL-8, IL-10, IL-12p70, according to the manufacturer’s instructions with some modifications. Briefly, standard curves with known concentrations of each cytokine were generated through double serial dilutions, resulting in an eight-point curve performed in duplicate. Five microliters of the supernatant were combined with a fluorescent bead’s reagent at a 1:1 volume ratio and incubated for two hours at room temperature. Following incubation, the beads were washed by resuspending them in wash buffer and then centrifuged (250 x g, 5 minutes). The supernatant was carefully discarded, and a mix of secondary antibodies was added and incubated for one hour. Subsequently, Streptavidin-Phycoerythrin (SA-PE) conjugates were introduced into the same reaction and incubated for 30 minutes at room temperature. After an additional washing step, the samples were analysed using a BD® FACSCelesta flow cytometer fitted with 405 nm, 488 nm, and 633 nm lasers and operated through the BD® FACSDiva software v.8. The detection limits for cytokines are IL-1β, 1.5 ± 0.6 pg/mL; TNF-α, 0.9 ± 0.8 pg/mL; IL-6, 1.5 ± 0.7 pg/mL; IL-8, 2.0 ± 0.5 pg/mL; IL-10, 2.0 ± 0.5 pg/mL; IL-12p70, 2.0 ± 0.2 pg/mL. Data analysis was performed using BD software, which is accessible online at LEGENDplex™ Cloud-based Data Analysis Software (https://legendplex.qognit.com).

### Genomic DNA extraction and 16S sequencing

Genomic DNA extraction from 48 colostrum samples was performed following an optimized phenol-chloroform protocol, as previously described.^13^ Briefly, 1 mL of colostrum was utilized for DNA extraction whenever feasible. The procedure involved the removal of the fat layer using a sterile hyssop and pelleting bacterial cells with a high-speed centrifugation followed by a sterile PBS wash. The resulting pellet was then treated with lysis buffer and mechanically lysed using a FastPrep system (MP Biomedicals, Santa Ana, CA). Enzymatic lysis with proteinase K, lysozyme, and RNAse was subsequently performed. DNA extraction was carried out using a phenol:chloroform:isoamyl-alcohol (25:24:1) mixture, followed by precipitation of DNA with isopropanol. The DNA pellet was washed with 70% ethanol and resuspended in nuclease-free water. As controls, an in-house mock with two bacterial isolates (*Escherichia coli* and *Pseudomonas putida* in a 1:1 ratio) and a DNA extraction of sterile PBS were included as positive and negative controls, respectively. DNA integrity was assessed by agarose gel electrophoresis, and concentration was determined using a NanoDrop ND-1000 UV spectrophotometer (Thermo Fisher Scientific, Waltham, MA, USA). Unless stated otherwise, all reagents were obtained from Sigma Aldrich.

For sequencing, the prepared DNA samples were subjected to Zymo Research’s Quick-16S kit, utilizing phased primers 341F (5’-CCTACGGGDGGCWGCAG-3’) and 805R (5’-GACTACHVGGGTATCTAATCC-3’) targeting the V3-V4 regions of the 16S rDNA. The sequencing was performed on a V3 MiSeq 622cyc flowcell, generating 2x301 bp paired-end reads.

### Colostrum taxonomic and differential profile

The sequencing reads were imported into QIIME2 v.2022.8 and demultiplexed for subsequent analysis.^21^ DADA2 plugin was used for denoising and quality control with optimized parameters to remove low quality regions of the sequences (--p-trim-left-f 12 --p-trim-left-r 16 --p-trunc-len-f 285 --p-trunc-len-r 250).^22^ Taxonomic species profiling was accomplished by aligning Amplicon Sequence Variants (ASVs) against the Silva 138.1 database with 99% of identity threshold^23^, using the q2-feature-classifier classify-sklearn naïve Bayes taxonomy classifier.^24^ To ensure data quality, potential contaminants identified through batch effect analysis or the positive control (Archaea, mitochondria/chloroplast, *Cyanobacteria*, *Streptomyces*, *Stenotrophomonas*, and *Pseudomonas*) were removed.^25^ Additionally, rare taxa were identified as ASVs with a total read count of ≤18 in at least three samples and subsequently excluded from the analysis.^26^

Rarefaction curves were generated to assess the sequencing depth and ensure adequate coverage of microbial diversity in colostrum. The rarefaction was conducted at a depth of 7,712 sequences. Diversity analyses were performed in R using vegan v2.6-4^27^ and phyloseq v1.44.0 libraries.^28^ Alpha diversity analyses included the calculation of Shannon diversity index, observed ASVs, evenness, and phylogenetic distance. Comparisons between groups were conducted using the Mann-Whitney’s U test. Beta diversity was assessed through ANOSIM (performed with 999 permutations) using the UniFrac and robust Aitchison distance matrices. A Principal Coordinates Analysis (PCoA) and a Robust Principal Component Analysis (RPCA) were created and visualized using EMPEROR.^29^ Differential abundance analysis at the taxonomic family level was conducted using DESeq2 v1.40.2 with batch correction applied^30^, and ANCOMBC v2.6.0.^31^ Only those families that were significantly different in both tools were graphed.

### Colostrum metabolite extraction

Due to limited sample volume availability, 25 colostrum samples (NW = 5 and WO = 20) were used to extract metabolites. 100 µL of colostrum was combined with 600 µL of acetonitrile (ACN). The mixture was vortexed for 1 minute and subjected to 30 minutes of sonication in ice-cold water to enhance the extraction of metabolites. Subsequently, the samples were centrifuged using optimized centrifuge parameters (20,817 x g, 4 °C for 10 minutes). From the resulting supernatant, 400 µL were carefully collected for vacuum evaporation of ACN. The dried metabolome was then resuspended in 100 µL of an injection solution composed of a water:acetonitrile mixture in an 80:20 ratio containing 0.1% formic acid (v/v). To ensure effective resuspension, the samples were vortexed for 1 minute and sonicated in ice-cold water for 30 minutes. To obtain a particle-free solution suitable for injection, the resuspended samples underwent centrifugation at the optimized parameters before being transferred to the injection vials. Quality control (QC) samples were generated by pooling all resuspended samples in equal volumes.

### LC-MS2 data acquisition

We employed the instrumentation and followed the equipment’s parameters previously described^32^, with minor modifications. Four μL of resuspended metabolomes were injected into an LC Agilent 1260 system (Agilent Technologies, Inc., Santa Clara, CA, USA). The separation of metabolites was done using a ProtID-Chip-43 II column (C18, 43 mm, 300 Å, 5 μm particle size, equipped with a 40 nL enrichment column). Mobile phase consisted of water with 0.1% formic acid (solution A) and acetonitrile (ACN) with 0.1% formic acid (solution B). The chromatographic method separation started with a mobile phase composition of 5% B, which was then linearly increased to 40% B over 20 minutes. Subsequently, the gradient was elevated to 100% B within 5 minutes and maintained at this composition for an additional 5 minutes. Next, the system was returned to its initial condition of 5% B within 1 minute and held for 9 minutes to ensure complete column re-equilibration before the next sample analysis. The total process time was 40 min, utilizing a flow rate of 300 nL/min. To mitigate potential carryover effects, two blank samples of 6 μL each - comprising one of injection solution and the other of ACN-were run between every sample injection. The separated metabolites were then introduced into an Agilent 6530A Q-TOF mass spectrometer (Agilent Technologies, Inc., Santa Clara, CA, USA) through a Chip Cube-LC interface using nano-spray ionization in positive mode. Data-dependent acquisition was used. For MS1, mass range was 110-2000 m/z with a 4 spectra/s velocity. The top 5 most intense precursor ions per cycle reaching 150 cps were selected for fragmentation (MS2 acquisition) in a mass range of 50-2000 m/z, at a 3 spectra/s rate. The active exclusion option was on, set to 2 spectra, and released after 0.25 min. Ramped collision energy was used with slope of 6 and offset of 4. Calibration was done before sample acquisition and every 24 hours with an ESI-L low mix concentration tuning mix solution (Agilent Technologies, Inc., Santa Clara, CA, USA) to ensure a mass accuracy <5 ppm for MS1 and MS2 data. The samples were randomly allocated in the autosampler for data acquisition.

### LC-MS2 data processing

Processing of LC-MS2 datasets was performed to reach two main goals: feature or peak (signals with *m/z* and retention time) extraction (peak-picking) and metabolite annotation at the structure (metabolomics standard initiative (MSI) levels 2 and 3), substructure, and chemical class (MSI, level 3) levels.^33^ Raw datasets were transformed from commercial .d format to open source .mzXML format using MSconvert tool within ProteoWizard v3.^34^ Transformed datasets were processed in MZmine v2.53 for peak-picking using the parameters described in Table S6.^35^ For feature annotation by spectral matching (MSI, level 2), open-source raw and processed data were uploaded and analyzed in the Global Natural Products Social Molecular Networking (GNPS) platform^36^ for classical molecular networking^37^ and feature-based molecular networking (FBMN)^38^, respectively. To expand the annotations not achieved by spectral matching, we employed MolDiscovery^39^, Dereplicator+^40^ within the GNPS environment. Additionally, we employed SIRIUS v5.8.1^41^ for database-independent chemical formula annotation and correction by ZODIAC^42^, in-silico structural metabolite annotation by CSI:FingerID^43^, and chemical class assignment by CANOPUS algorithm (MSI, level 3). The chemical landscape representation was created using the molecular network provided by the FBMN analysis in GNPS, enriched with the chemical superclass annotations of CANOPUS. Substructure (motifs; MSI, level 3) annotation was done using the MS2LDA webpage.^44^ Molecular structures assignments were filtered by a mass error less than 10 ppm to keep annotations with high-mass accuracy. Additionally, peaks and annotations derived from blank samples were filtered out from the analysis.

### Quantification and statistical analysis

Unless other specified, all statistical analyses were performed and plotted using GraphPad Prism version 8.0.1, GraphPad Software, San Diego, California USA (www.graphpad.com). The clinical data (maternal age, gestation weeks, delivery mode and neonatal sex) were compared between groups using independent-samples median test or Fisher’s exact test to evaluate their influence as confounding factors. Kruskal-Wallis test and GLM were used to assess the influence of maternal and neonatal demographic and clinical data on the microbiota composition, metabolomic profile, and cytokine quantification. Mann-Whitneýs U test was used for evaluating the difference in cytokine concentrations between the NW and WO groups. A p-value ≤ 0.05 was considered statistically significant.

For metabolomics data analysis, feature abundance normalization was conducted using the quantile method, and subsequent differential abundance analysis was carried out within the NormalyzerDE online platform.^45^ Metabolites exhibiting a log2-fold change of ±0.58 and a p-value <0.05 (calculated with the limma package^46^) were considered differentially abundant between WO and NW groups. Volcano plots were created to facilitate data visualization with the EnhancedVolcano R package v1.16.0.^47^ To assess differences among groups at the chemical class level, the summed abundance (area under the curve) of all the quantified features belonging to each respective class was calculated followed by a two-group comparison using the Wilcoxon test, assuming unequal variance, implemented with the wilcox.test() function in R. For comparing colostrum chemical diversity between groups, the Shannon and Simpson indexes were calculated as community ecology metrics^48^ using the vegan package v2.6-4. Statistical significance was determined through a two-group comparison using the Wilcoxon test, as described above.

Bacterial interactive network was constructed by linking taxonomic families that exhibited a Spearman rank correlation exceeding an absolute value of 0.5 and a p-value < 0.05. Network analysis was performed using MicrobiomeAnalyst.^49^ Correlations between clinical data, cytokine concentrations, bacterial families, and metabolite abundances were calculated with a Spearman’s correlation (ρ) using R. Only correlations with a p < 0.10 were kept for visualization in the heatmaps.

## Results

### Demographic and clinical characteristics of the study cohort

This study cohort consisted of 48 women between 20 and 32 years old. As per study design body mass index (BMI) was significantly higher in the group “With Obesity” (WO) compared to “Normal Weight” (NW) group (medians: 22.45 vs. 32.05, in NW and WO, respectively, p<0.0001, Table 1). There was no difference in maternal age, duration of gestation, delivery mode or biological sex of the offspring.

**Table 1.**
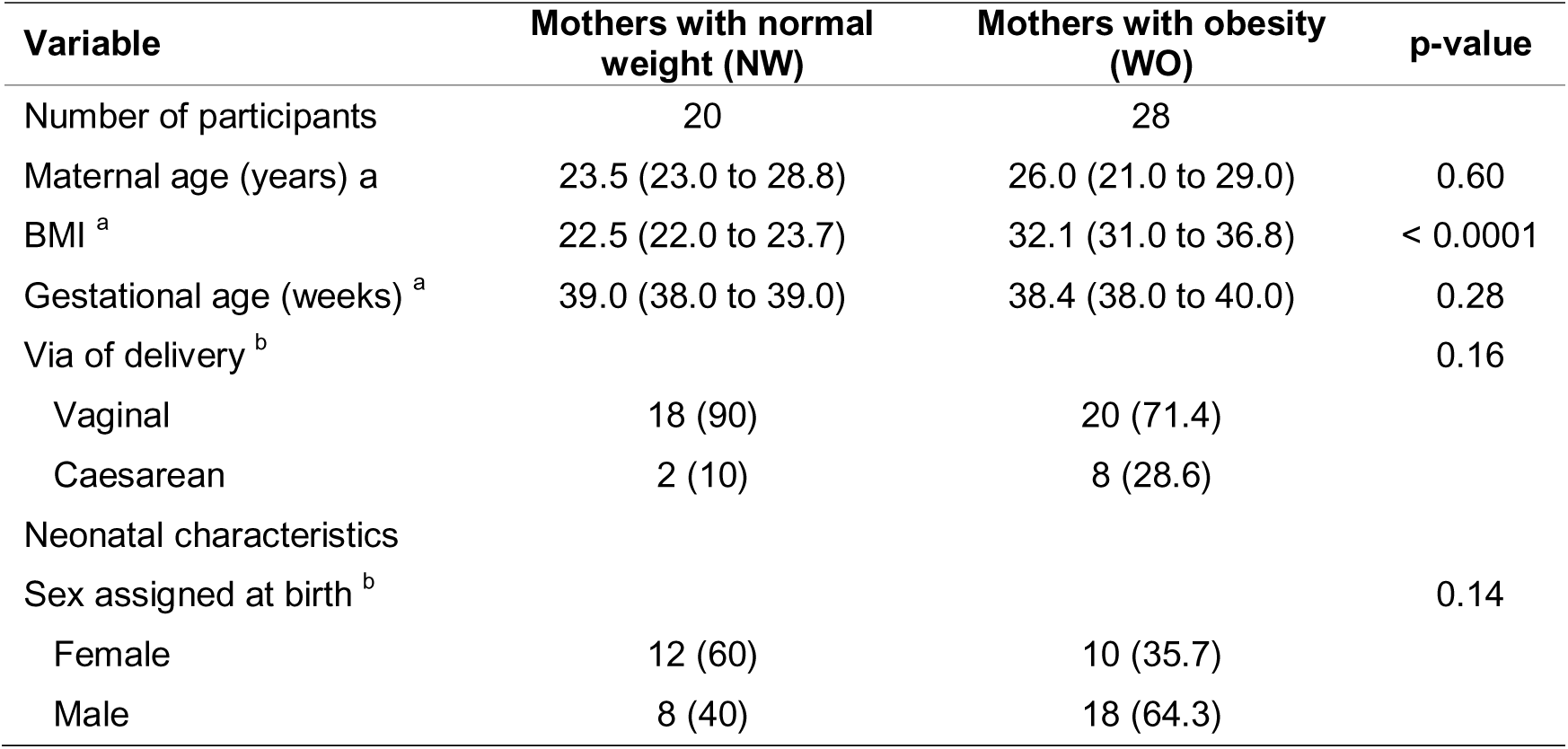
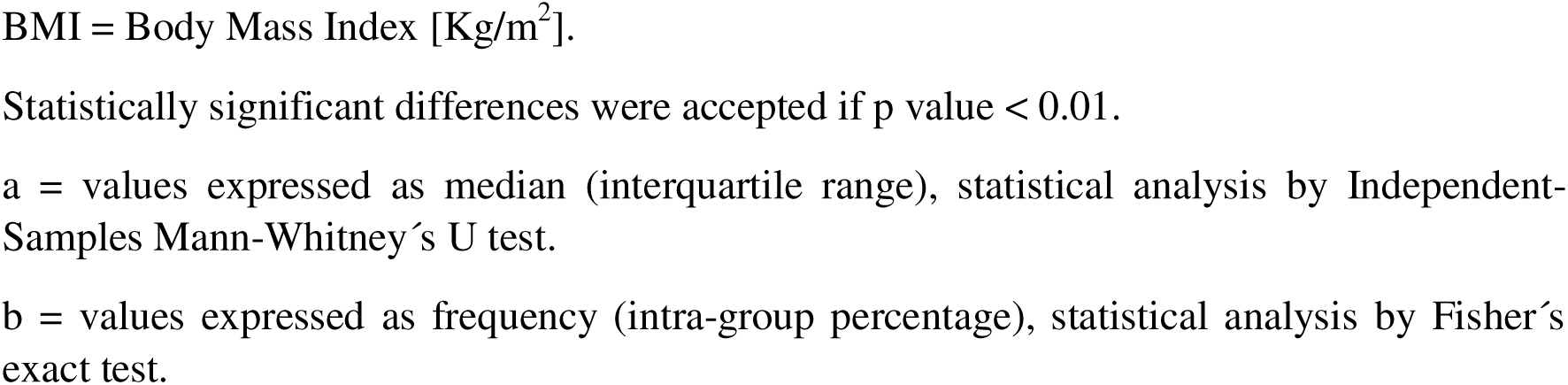
Study population characteristics, grouped in accordance with maternal weight classification.

| Variable | Mothers with normal weight (NW) | Mothers with obesity (WO) | p-value |
| --- | --- | --- | --- |
| Number of participants | 20 | 28 |  |
| Maternal age (years) <sup>a</sup> | 23.5 (23.0 to 28.8) | 26.0 (21.0 to 29.0) | 0.60 |
| BMI <sup>a</sup> | 22.5 (22.0 to 23.7) | 32.1 (31.0 to 36.8) | < 0.0001 |
| Gestational age (weeks) <sup>a</sup> | 39.0 (38.0 to 39.0) | 38.4 (38.0 to 40.0) | 0.28 |
| Via of delivery <sup>b</sup> |  |  | 0.16 |
| Vaginal | 18 (90) | 20 (71.4) |  |
| Caesarean | 2 (10) | 8 (28.6) |  |
| Neonatal characteristics |  |  |  |
| Sex assigned at birth <sup>b</sup> |  |  | 0.14 |
| Female | 12 (60) | 10 (35.7) |  |
| Male | 8 (40) | 18 (64.3) |  |

BMI = Body Mass Index [Kg/m<sup>2</sup>].
Statistically significant differences were accepted if p value < 0.01.
a = values expressed as median (interquartile range), statistical analysis by Independent-Samples Mann-Whitney's U test.
b = values expressed as frequency (intra-group percentage), statistical analysis by Fisher's exact test.

### There are no evident alterations in colostrum proinflammatory cytokines in maternal obesity

We quantified colostrum cytokines relevant in inflammatory processes. All cytokines were detected in all samples. However, no significant differences were observed between groups nor correlated with maternal BMI (Table 2 and Figure S1). Investigating relationships between maternal clinical characteristics and colostrum cytokine concentrations, evidenced that only maternal age presented general trends whereby increasing age led to decreasing cytokine concentrations for NW, except IL-1β, while the opposite trends were seen for mothers with obesity (Figure S2). In the WO group, colostrum TNF-α concentrations were negatively correlated with age (ρ = -0.43) (Figure S2).

**Table 2.**
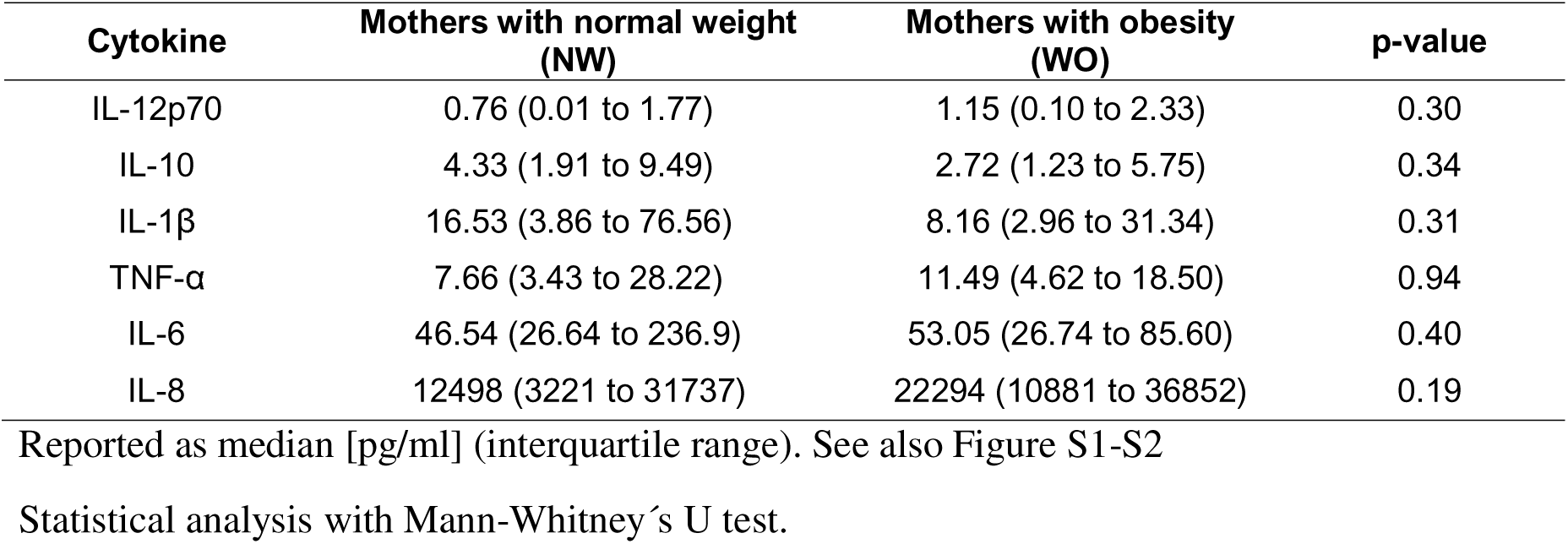
Concentrations of cytokines in colostrum.

### Changes in microbial composition in the colostrum of mothers with obesity correlates with previous findings

We analysed colostrum microbial community using 16S ribosomal RNA amplicon sequencing. We obtained 2,280,755 high-quality reads, corresponding to 45,615 reads per sample. We identified 1,255 amplicon sequence variants (ASVs) annotated within 4 major phyla: Firmicutes (66.5%, NW; 77.5%, WO), Proteobacteria (21.9%, NW; 12.3%, WO), Actinobacteriota (4.5%, NW; 4.8%, WO), and Bacteroidota (2.3%, NW; 1.6%, WO) (Figure 1A). Overall, the taxonomic families *Staphylococcaceae* (37.8%, rank 0.17%-97.8%) and *Streptococcaceae* (28.8%, rank 0.02%-92.7%) exhibited the highest relative abundance. Lower abundances were observed for *Gemellaceae* (3.7%), *Burkholderiaceae* (2.2%), *Enterobacteriaceae* (2%), and *Sphingomonadaceae* (1.3%) taxonomic families. Additional families (102), including *Bacillaceae*, *Lactobacillaceae*, *Bifidobacteriaceae*, contributed to <1% of the total relative abundance (Figure 1B, Table S1). There was no difference between groups looking at the main ecological diversity indices (Figure S3). The Firmicutes to Bacteroidota (F/B) ratio and the Firmicutes to Proteobacteria (F/P) ratio were higher in samples from women with obesity, although these differences were not statistically significant (mean ratio 28.7 vs 47.7, p = 0.30, and 3.0 vs 6.3, p = 0.14, respectively). In contrast, the Proteobacteria to Bacteroidota (P/B) ratio was significantly decreased in WO (9.4 vs 7.6, p = 0.046), which agrees with a relative reduction in the abundance of the Proteobacteria phylum, although not significant (21.9% ± 23.5% vs 12.3% ± 17.2%, p = 0.14).

**Figure 1.**
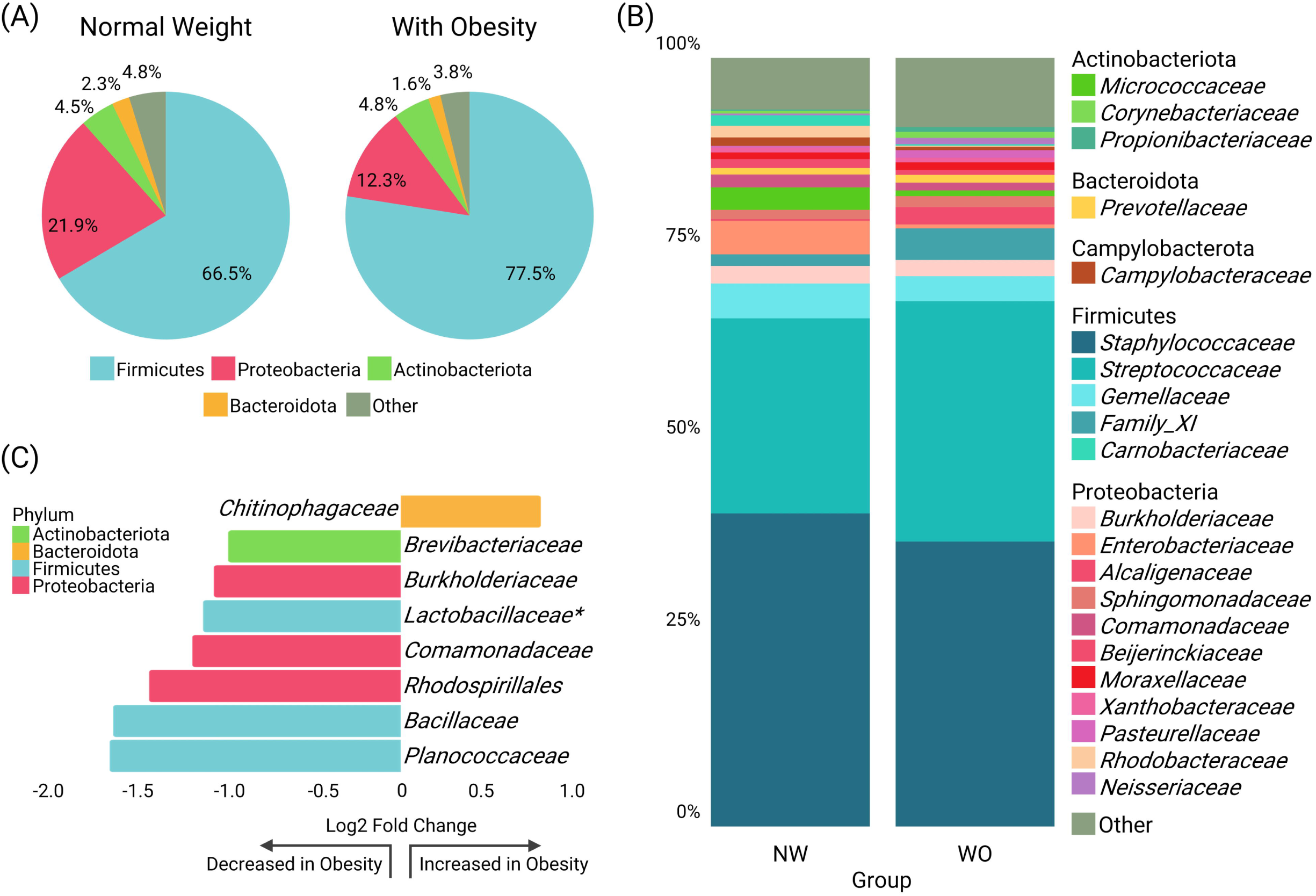
Composition of the microbiota of colostrum collected from parent with and without obesity. (A) Relative abundance of the most abundant taxa at phylum and (B) family level. “Other” classification represents bacterial families with less than 1% of the total relative abundance. (C) Differential abundance analysis performed at the taxonomic family level using ANCOMBC. The graph provides information on the phylum, family, and corresponding Log2 Fold Change values (p <0.05), which indicate the magnitude and direction of the differential abundance. NW, normal weight group (n=20); WO, with obesity group (n=28); *p <0.10. See also Table S1.

In the WO group, the *Chitinophagaceae* family was 0.78-fold increased (Figure 1C). In contrast, 7 taxonomic families were less abundant, including *Brevibacteraceae* (Log2FC - 0.96), *Burkholderiaceae* (Log2FC -1.04), *Lactobacillaceae* (Log2FC -1.10), *Comamonadaceae* (Log2FC -1.16), *Rhodospirillales* (Log2FC -1.40), *Bacillaceae* (Log2FC - 1.60), and *Planococcaceae* (Log2FC -1.62).

### Maternal obesity correlates with changes in the colostrum metabolite environment

We used untargeted metabolomics to explore discrepancies in the colostrum of women with obesity compared to normal weight subjects. Our pipeline led to the quantification of 2,808 features across samples. We annotated 1,027 MS2-containing features across 12 superclasses, leaving 473 features unassigned (Figure 2A, Table S2). Metabolite diversity at the class level was assessed using the Shannon and Simpson indexes, revealing a reduction in colostrum samples from WO compared to NW (p = 0.05; Figure S4). Subsequently, we observed 73 features with decreased abundance and 49 features with increased abundance in the WO group (log2FC ± 0.58, limma test p < 0.05; Figure 2B). Decreased features were mostly linked to carboxylic acids, fatty acyls and prenol lipids, whereas the distribution of the increased features was primarily carboxylic acids, organooxygen compounds, azoles, and organic phosphines (Figure 2C). To gain insights into metabolic markers of obesity, we performed correlation analyses between metabolites (identified at the chemical class level) and maternal BMI levels. We identified 57 features distributed across 11 chemical classes that were significantly associated with maternal BMI (Figure 2D and Table S3), mainly comprising carboxylic acids and derivatives (n = 31), followed by fatty acyls (n = 7), and organooxygen compounds (n = 6). Through high-confidence putative structural annotation (mass error ≤ 10 ppm) and differential abundance analysis (limma test), we found that decanoylcarnitine (alongside an unidentified carnitine, as revealed through MS2LDA analysis) and 15-HEDE (fatty acyl) presented reduced abundance in the colostrum of WO compared to NW group (Figure 2E and Table S4).

**Figure 2.**
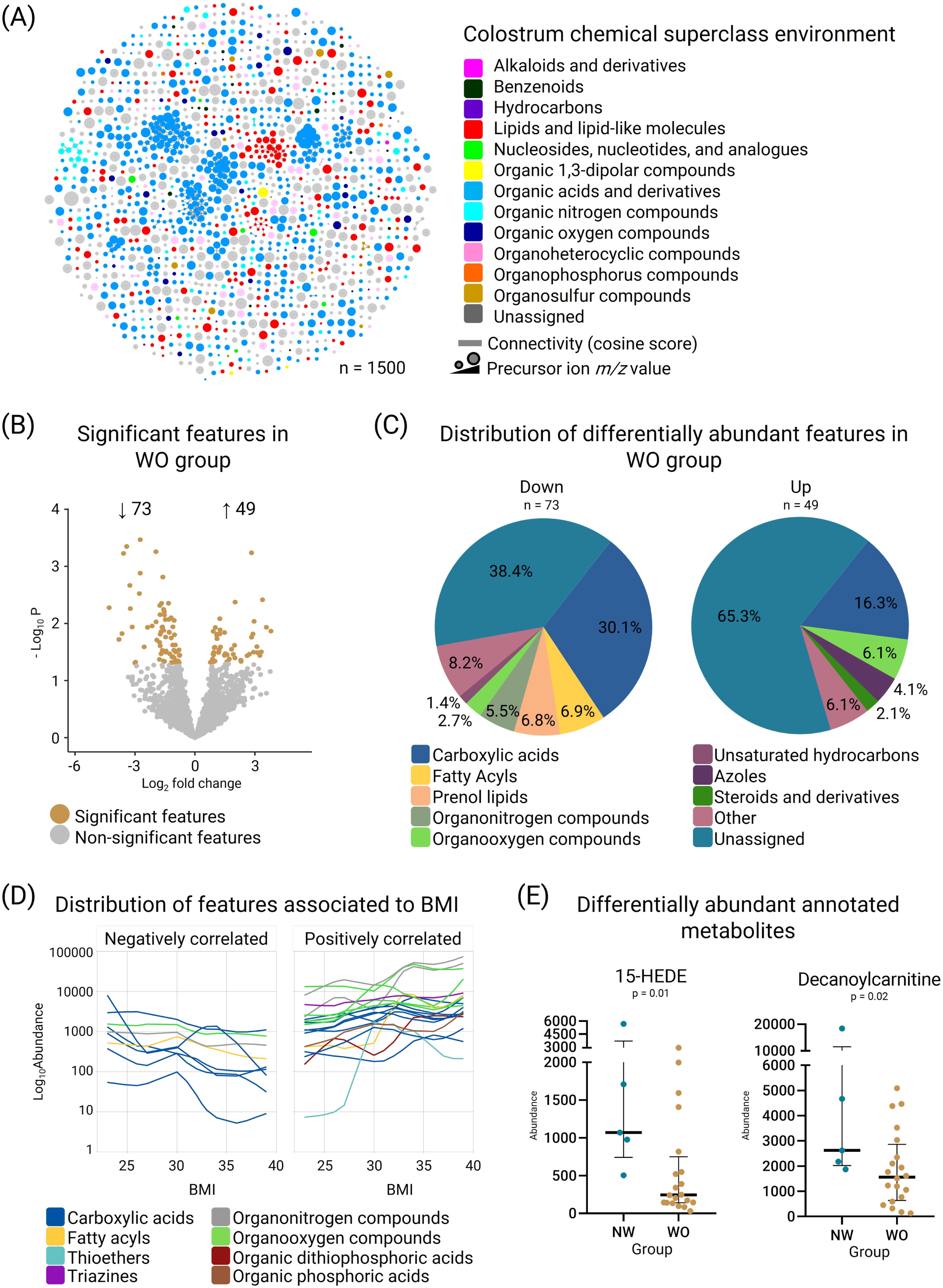
Metabolite environment in colostrum from women with obesity (WO). (A) Molecular network of the identified MS2-containing features in colostrum samples. Each node represents a feature, and the color denotes its assigned chemical superclass. Node size highlights the m/z value. Edge thickness indicates the MS2 similarity (cosine score) among features; (B) Volcano plot of all quantified features highlighting the differentially abundant features between WO/NW groups. Brown dots represent significantly increased or decreased features in the WO group (log2FC ± 0.58, limma p < 0.05), while gray dots indicate features without significant differences; (C) Pie charts of the chemical class distribution of the differentially increased (n=49) or decreased (n=73) features in colostrum samples from the WO group. “Other” classification corresponds to classes that only contained one feature; (D) Loess curves of each feature significantly associated with maternal BMI classified by their Spearman’s correlation value (ρ). Curves are color-coded based on the feature chemical class assigned as described in the color key. Only features with a correlation value higher than absolute 0.45 were plotted; (E) Boxplots of the relative abundance of the dysregulated (limma test, p < 0.05) lipid related identified metabolites, 15-HEDE and decanoylcarnitine; NW, normal weight group (n=5); WO, with obesity group (n=20). See also Table S2-S4.

### Bacterial co-occurrence network in colostrum microbiota

Expanding upon the abundance analysis of individual elements, we used co-occurrence networks to define distinctive bacterial interaction modules within the samples. This approach provides a comprehensive view of the coexistence of bacterial taxa and their potential influence with each other. Using Spearman rank coefficients, we defined five distinct modules of taxonomic families according to their degree of interaction. A positive correlation in this context denotes a potential coordinated and mutualistic interaction, either through direct or indirect mechanisms, which implies that the presence of specific taxa positively influences the presence of others within the microbial community. Conversely, a negative correlation indicates a direct or indirect antagonistic interaction, wherein the coexistence of certain taxa is perturbed by the presence of others (Figure 3 and Table S5).

**Figure 3.**
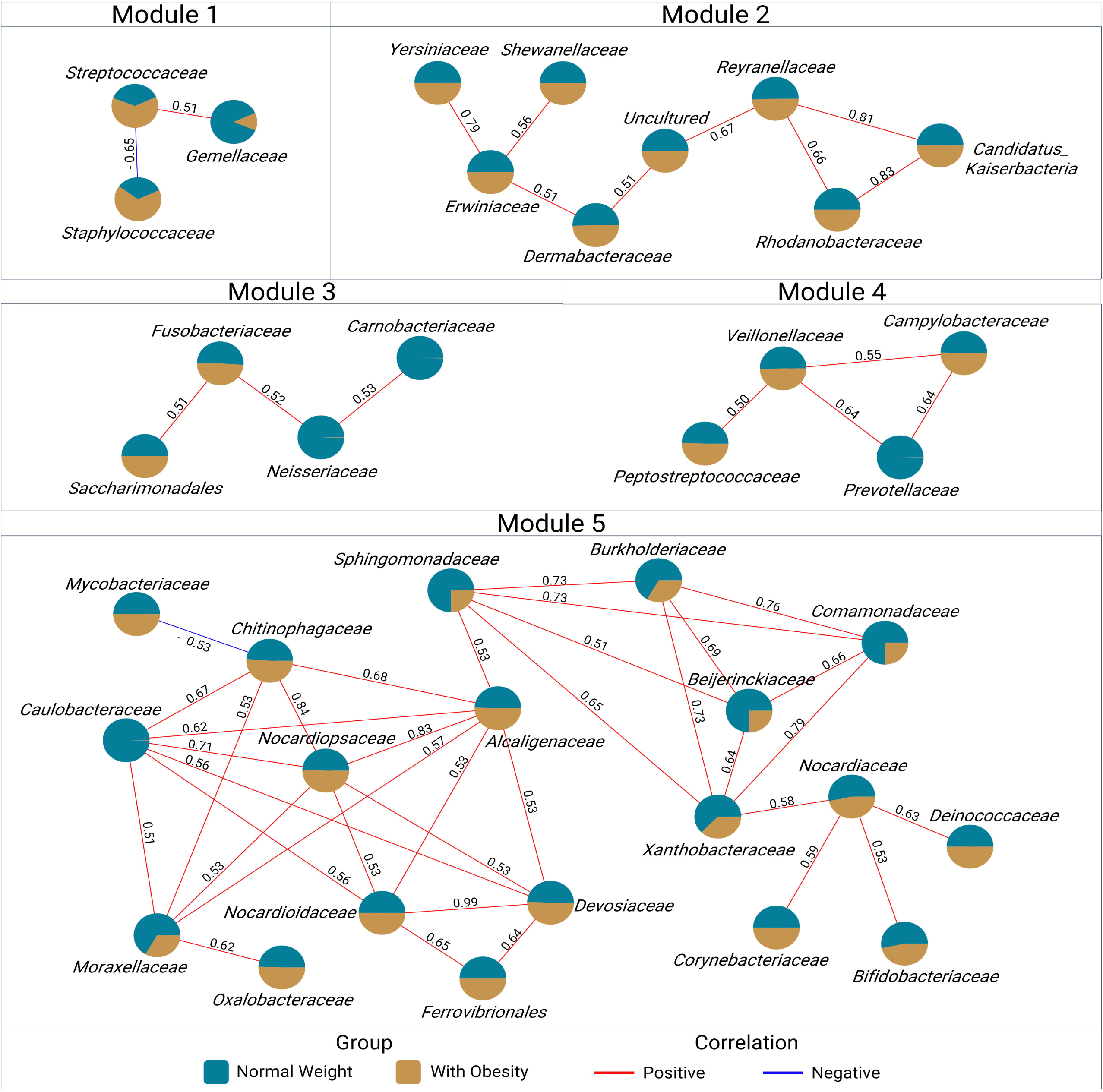
Colostrum bacterial co-occurrence network. Visualization of the interactions between taxonomic families’ abundances within the normal weight (blue; n=20) and with obesity (brown; n=28) groups. Five distinct modules, each comprising at least 3 members, were obtained based on Spearman’s rank correlation of coexistence. Each node is represented by a pie chart displaying the median relative abundance of a taxonomic family within each group. Edges connect two families if their Spearman’s rank correlation exceeds an absolute value of ρ = 0.5 with a p-value < 0.05. The edges are annotated with Spearman’s correlation values. Positive correlations between differentially abundant taxonomic families are represented by red edges, while negative correlations are indicated by blue edges as described in the color key. See also Table S5.

In the first module, we observed relationships among Firmicutes members, with *Streptococcaceae* negatively correlated to *Staphylococcaceae* but positively correlated to *Gemellaceae*. The second module includes mostly members of Proteobacteria positively correlated, including *Yersiniaceae*, *Erwiniaceae*, *Shewanellaceae*, *Reyranellaceae* and *Rhodanobacteraceae*. In the third module we identified *Saccharimonadales*, *Fusobacteriaceae*, *Neisseriaceae* and *Carnobacteriaceae,* positively correlated. The fourth module presented positive correlations between *Prevotellaceae*, *Campylobacteraceae*, *Veillonellaceae* and *Peptostreptococcaceae*. Finally, the fifth module, includes 19 taxonomic families with positive correlations. Notably, in this module, we identified families with reductions in relative abundance in the colostrum of women with obesity, including *Comamonadaceae*, *Burkholderiaceae*, *Xanthobacteraceae*, *Beijerinckiaceae*, *Sphingomonadaceae*, *Caulobacteraceae*, *Bifidobacteriaceae*, *Nocardiaceae*, *Deinococcaceae*, and *Moraxellaceae*. Further investigation is required to unravel the complex network of interactions that may hold biological significance in the context of maternal obesity.

### Lipid mediators are correlated with proinflammatory cytokines and proteobacteria members in colostrum

We performed co-occurrence analysis to explore potential interactions between putatively annotated metabolites and cytokines in colostrum (Figure 4A-B). We found that 15-HEDE negatively correlated with IL-12p70 and IL-10, suggesting a potential role of this regulatory lipid in inflammation. Lysophosphatidylcholine (16:00; LysoPC [16:00]), on the other hand, correlated positively with IL-1 and IL-8, suggesting a role in proinflammatory signalling pathways.

**Figure 4.**
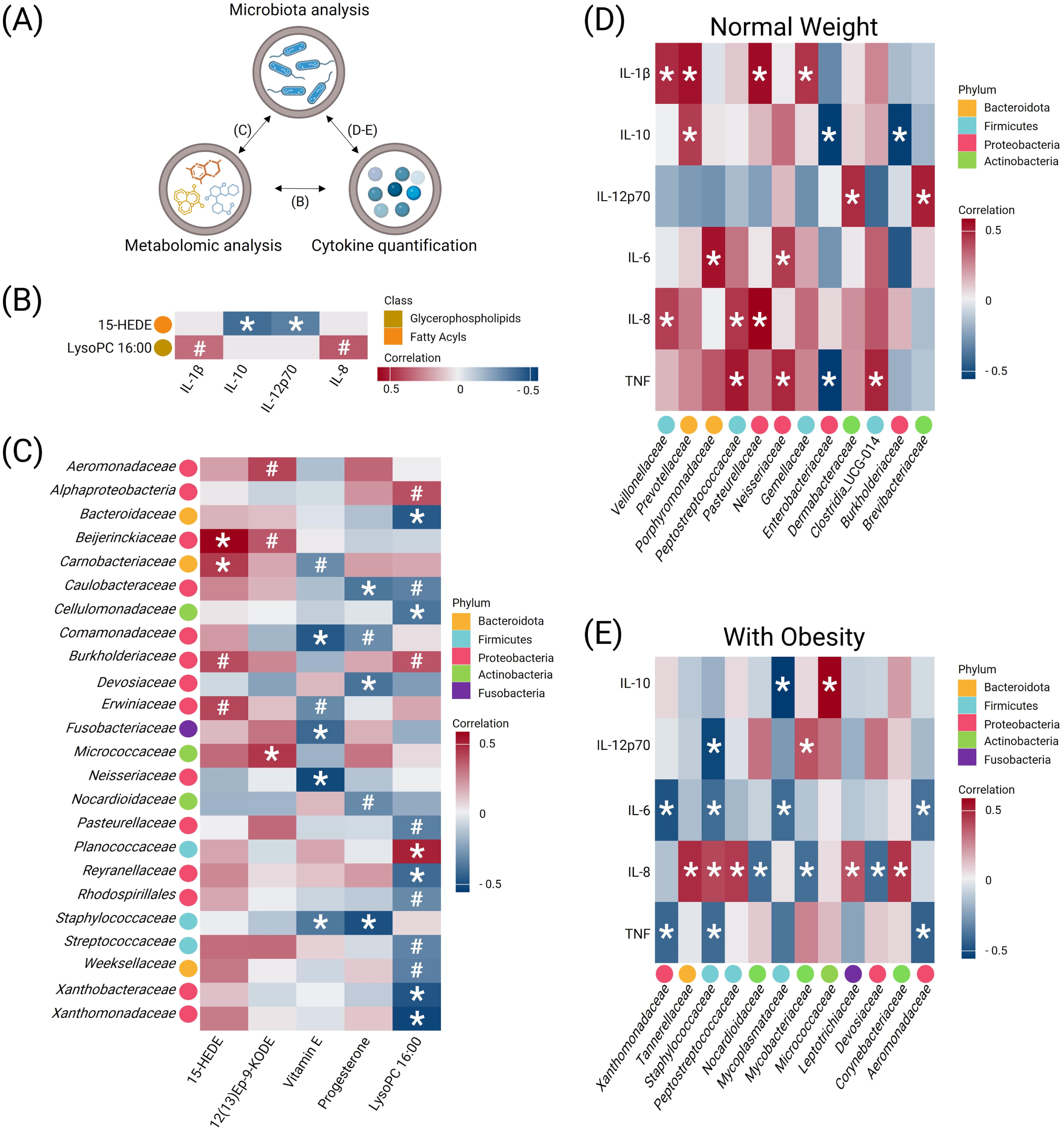
Correlation analysis between colostrum bioactive elements. (A) Scheme of the different correlations performed between the bioactive elements. (B) Heatmap of the Spearman’s correlation between metabolites and cytokines in colostrum; (C) Heatmap of the Spearman’s correlation analysis of taxonomic families and metabolites in colostrum; (D-E) Heatmap of the Spearman’s correlation analysis of taxonomic families and cytokines in colostrum from (D) normal weight and (E) with obesity women. Red represents positive correlation, white represents low correlation, and blue represents negative correlation, as shown in the color key. *p-value < 0.05; #p-value < 0.10.

In addition, we analysed the interrelationship between metabolites and bacterial groups. For instance, the regulatory lipid 15-HEDE, found to be decreased in colostrum of women with obesity, exhibited positive correlations with taxonomic families such as *Beijerinckiaceae*, *Burkholderiaceae*, and *Erwiniaceae*, which were also less abundant in colostrum of women with obesity (Figure 1C and Figure 4C). Moreover, despite the absence of a significant difference between study groups, 12(13) Ep-9-KODE showed a positive correlation with *Micrococcaceae*, *Aeromonadaceae*, and *Beijerinckiaceae*. On the other hand, LysoPC (16:00) showed negative correlations with *Xanthobacteraceae*, *Reyranellaceae*, *Xanthomonadaceae*, *Caulobacteraceae*, *Cellulomonadaceae*, *Pasteurellaceae*, *Rhodospirillales*, *Streptococcaceae*, *Weeksellaceae* and *Bacteroidaceae,* but exhibited positive correlations with *Planococcaceae*, *Alphaproteobacteria* and *Burkholderiaceae*.

We also identified that the steroid hormone progesterone displayed negative correlations with *Caulobacteraceae*, *Comamonadaceae*, *Devosiaceae*, *Nocardioidaceae* and *Staphylococcaceae,* which may imply instances where negatively affects abundance of the bacterial community. Furthermore, fat-soluble vitamin E was negatively correlated with *Comamonadaceae*, *Neisseriaceae*, *Fusobacteriaceae*, *Staphylococcaceae*, *Erwiniaceae* and *Carnobacteriaceae* (Figure 4C). Further research into the specific metabolic pathways and ecological roles associated with these metabolites could support their significance in the colostrum system and their implications in obesity.

### Proinflammatory cytokines are negatively correlated to Proteobacteria members in colostrum of mothers with obesity

We also delve into the interplay between microbial communities and immune-modulating factors. In samples obtained from normal weight participants, members of the Proteobacteria phylum such as *Enterobacteriaceae* and *Burkholderiaceae* displayed negative correlations with IL-10, while *Prevotellaceae* showed a positive correlation with this anti-inflammatory cytokine. Other members of the Proteobacteria phylum, including *Neisseriaceae* and *Pasteurellaceae*, exhibited positive correlations with IL-6, TNF-α, IL-1β, and IL-8, suggesting a potential proinflammatory modulation (Figure 4D). Meanwhile *Veillonellaceae*, *Prevotellaceae*, and *Gemellaceae* presented a positive correlation with IL-1β, *Dermabacteraceae* and *Brevibacteraceae* were positively correlated to IL-12p70. Additionally, *Porphyromonadaceae* was positively correlated to IL-6.

In contrast, samples from women with obesity showed negative correlations between the taxonomic families *Aeromonadaceae*, *Xanthomonadaceae*, and *Staphylococcaceae*, and the immune-modulating factors IL-6, TNF-α, and IL-12p70. *Devosiaceae*, *Mycobacteriaceae*, and *Nocardioidaceae* were negatively correlated with IL-8, while *Corynebacteriaceae*, *Staphylococcaceae*, *Leptotrichiaceae* and *Tannerellaceae* showed positive correlations (Figure 4E). Notably, *Peptostreptococcaceae* exhibited a positive correlation with IL-8 across the entire cohort independently of the study group.

## Discussion

Colostrum is a complex and dynamic fluid that seeds neonatal intestines within the first hours of life. The interactions among colostrum bioactive components are thought to significantly impact the structure and stability of its microenvironment.^10,14^ Research initiatives, such as those within the BEGIN project, are exploring breastmilk as a biological system, emphasizing the interactions, and coordinated functionality between its bioactive components.^10,50^ However, understanding how maternal obesity influences the bioactive composition of colostrum and the impact on neonatal immune system remains limited. In this study, we quantitatively profiled the colostrum microbiota, cytokines, and metabolites to establish networks of co-occurring elements, aiming to provide proof of concept for colostrum as a system and elucidate the effect of maternal obesity on shaping their associations.

Despite extensive research on the associations between breastmilk components and maternal obesity, the findings remain heterogeneous.^4,11,13,51–53^ We identified subtle differences while analysing individual bioactive components *i.e.* cytokines, microbiota, and metabolites. This results variability can be attributed to differences in study population, sample size, lactation stage, and methodological techniques applied.^54–56^ However, addressing colostrum as a dynamic biological system with interconnected components, our differential and co-abundance assessments provided valuable insights into the colostrum microbial ecology. Co-occurrence analysis elucidates potential relationships within microbial communities and their interactions with bioactive molecules. In natural settings, microorganisms interact (mutualism, competition, and commensalism) leading to the formation of complex networks characterized by non-random co-occurrence patterns.^19^ Co-occurrence analysis offers insights into microbial community structure that traditional methods may overlook, for example by simply characterizing microbial diversity.^57^ Understanding microbial associations within a co-occurrence network allows for the identification of key interactions for community organization and proper functioning of ecosystems.^58^ In our study, we identified a negative correlation between *Staphylococcaceae* and *Streptococcaceae* (ρ = -0.65), indicating a potential competitive relationship between these bacterial taxa. This association may be influenced by ecological factors within the mammary gland microenvironment, such as niche competition, resource availability, or immune responses, which could benefit the dominance of one bacterial group over the other.^59,60^ *Staphylococcus*/*Streptococcus* competition in breastmilk, with infants fed *Staphylococcus*-dominated breastmilk exhibiting a more complex gut and nasal microbiota networks than those fed *Streptococcus*-abundant breastmilk during early infancy.^14^ Further investigation is warranted to elucidate the underlying mechanisms driving these predominant bacterial dynamics in colostrum and their implications for maternal and infant health.

Moreover, co-occurrence network analysis revealed bacterial relationships among Proteobacteria members exhibiting reduced relative abundance in colostrum from women with obesity. Within the same co-occurrence module, these taxa demonstrated positive correlations, indicating mutualistic interactions and functional similarities.^59^

The observed variations in colostrum microbiota from WO group may impact the microbiota development of neonates consuming this milk. Offspring born to mothers with obesity presented a decrease in the abundance of the early pioneering Gammaproteobacteria two weeks after birth.^61^ This is critical for the establishment of a healthy gut microbiome functioning since Proteobacteria members are the primary colonizers during the first week of life and play a crucial role in preparing anaerobic conditions in the gut for successive colonization by strict anaerobes.^62–64^ Additionally, their presence in the postnatal period is essential for promoting a healthy immune system, facilitating host-microbe interactions. Consequently, a reduction in Proteobacteria supply in colostrum from mothers with obesity may compromise immune system training during the critical early days of life, potentially leading to impaired immunological tolerance and increased risk of immune-related diseases later in life.^65,66^

Alterations in specific microbial taxa within breastmilk may potentially be linked to cytokine levels, exerting an impact on breast-fed infants.^67^ We report that abundance of *Enterobacteriaceae* in colostrum from women with normal weight was negatively correlated with TNF-α (ρ = -0.54), while Proteobacteria such as *Neisseriaceae* was positively correlated with TNF-α and IL-6 (ρ = 0.53 and ρ = 0.48, respectively). This pattern aligns with previous observations in mice, where the presence of the early colonizer *Enterobacteria* triggers TNF-α production during the first week of life. This interaction prevents excessive inflammatory and autoimmune gastrointestinal disorders in the future.^66^ Furthermore, the production of TNF-α by macrophages and monocytes within the first three days of life, stimulated by the microbiota, is a prerequisite for the maturation of pre-conventional dendritic cell 1 (pre-cDC1) into conventional dendritic cell 1 (cDC1).^68^

The increased levels of Gram-negative bacteria in the gut are known to stimulate the local production of proinflammatory cytokines.^69^ In our work, we provide information suggesting that such phenomenon may be occurring in the mammary gland. In mothers with obesity (WO group), we observed negative correlations between proinflammatory cytokines (TNF-α, IL-6, and IL-12p70) and certain bacterial families such as *Aeromonadaceae*, *Xanthomonadaceae* and *Staphylococcaceae* (Figure 5). Our unexpected correlative findings align with observations of lower TNF-α and IL-6 concentrations in breastmilk from mothers with obesity compared to mothers with normal BMI.^70^ This suggests a potential local regulatory system within the mammary gland that may modulate systemic inflammation during obesity.

**Figure 5.**
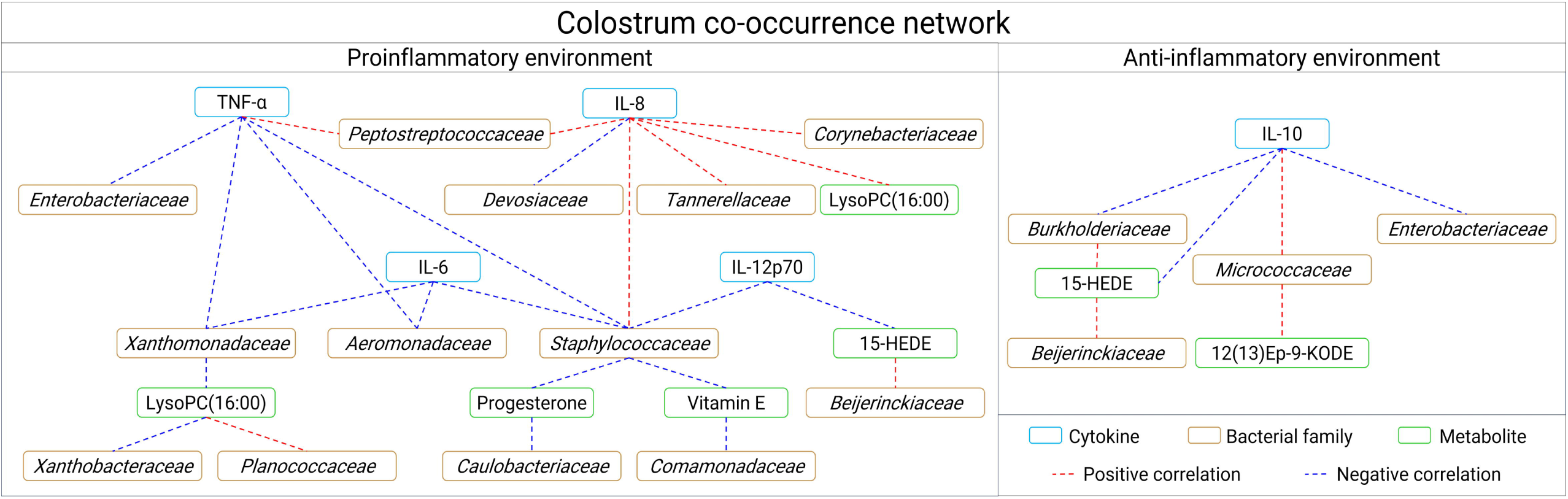
Integrated biological network of colostrum environment in obesity context. Summary of Spearman’s correlation analysis between colostrum inflammatory cytokines quantification, bacterial families, and metabolites in the maternal obesity context (p < 0.05). Left panel corresponds to correlation network between proinflammatory cytokines and bioactives in colostrum samples from women with obesity. Right panel corresponds to colostrum anti-inflammatory cytokines correlation network from women with obesity. Rectangle colors represent bioactive elements: blue rectangle corresponds to cytokines, brown to bacterial families, and green to metabolites. Red and blue edges represent the positive and negative correlation, respectively.

Comprehensive metabolite quantification is necessary to deeper analyze colostrum ecological system of women with obesity. These interactions have been associated with variations in breastmilk metabolites.^5,71,72^ In our cohort, we observed a reduction in fatty acyls and prenol lipids, which suggests potential alterations in the lipid profile of colostrum associated with maternal obesity. For instance, we observed that decanoylcarnitine exhibited a negative correlation (ρ = -0.57) with BMI and was found to be decreased in the WO group. Our results contrast previous observations where elevated levels of various acylcarnitines were reported in serum and breastmilk from individuals with obesity compared to non-obese and metabolically healthy obese individuals.^5,73^ Acylcarnitines play a crucial role in mediating the beta-oxidation of fatty acids within mitochondria, a pivotal process in energy production.^74^

In addition, breastmilk of women with overweight or obesity exhibit decreased levels of polyunsaturated fatty acids (PUFAs).^75^ The metabolism of PUFAs processed through the enzymatic action of lipoxygenases and cyclooxygenases can lead to the formation of oxylipins.^76^ Breastmilk has a diverse array of bioactive lipids, which play significant roles within the neonatal immune system.^77,78^ The presence of oxylipins in breastmilk may serve as a significant indicator of maternal health and could influence the health outcomes and inflammatory status of breast-fed infants.^76^ Some hydroxy fatty acids appear to be involved in complex and varied functions linked to inflammatory process with robust functions, for example with anti-inflammatory and pro-resolving properties, either directly or through their conversion into lipid mediators.^79,80^

One notable example is 15-HEDE, a macrophage-oxidized product of the inflammatory modulator 6-PUFA eicosadienoic acid (EDA)^81,82^, which in our study was found to be significantly decreased in samples from women with obesity and exhibited a negative correlation with IL-10 levels, while it was positively correlated with Proteobacteria members, such as *Beijerinckiaceae* and *Burkholderiaceae* (Figure 5). Despite that 15-HEDE has been described as an inhibitor of 5-lipoxygenase (5-LO)^83^, it has also been reported to induce of vasodilation and vascular hyperpermeability, acting as a proinflammatory lipid mediator that exacerbates allergic rhinitis.^84^ Elevated levels of 15-HEDE have also been observed in allergic airway inflammation in mice after a house dust mite exposure and in psoriasis.^81,85^ The negative correlation with IL-10 could suggests that IL-10 may act as a modulator of inflammation resolution in colostrum. In the chronic inflammatory milieu of obesity, circulating levels of IL-10 are elevated compared to normal weight women^86,87^, thereby we hypothesized a potential compensatory mechanism to counterbalance chronic inflammatory processes and mitigate the impact of reduced but highly immunogenic bacteria. In normal-weight mothers, the higher abundance of 15-HEDE may enhance the vasodilatory capacity of the mammary vasculature, facilitating efficient nutrient transfer and immunological component delivery to the neonate.^88^ This process likely supports optimal colostrum composition and functionality, contributing to the establishment of the neonatal gut microbiota and immune system. These results demonstrate the connection of altered lipidic metabolite concentrations, disrupted bacterial community, and the local inflammatory markers at colostrum environment within the context of maternal obesity, which may influence the neonatal colonization.

While progesterone showed no significant abundance difference between the NW and WO groups, we observed negative correlations when compared to specific bacterial taxa, including *Staphylocococcaceae*, *Ferrovibrionales*, *Caulobacteriaceae*, and *Devosiaceae* (Figure 5). Progesterone has been shown to influence microbial communities in vitro, by reducing the growth, adhesion, and virulence of certain *Staphylococcus* strains.^89^ Bacteria from Actinobacteria, Proteobacteria, and Firmicutes phyla express hydroxysteroid dehydrogenases (HSDs), key enzymes involved in steroid hormone catabolism, potentially using steroids as a carbon source.^90^ While progesterone is dominant during late pregnancy, pregnant women with obesity often exhibit reduced serum progesterone levels.^91,92^ Moreover, progesterone levels in breastmilk from women with a high fat and protein diet are diminished^93^ and negatively correlated with infant weight at 6 months.^94^

Our results provide valuable proof of concept into the systemic structure of colostrum and the impact of maternal obesity on its ecology. However, this study presents some limitations. For instance, sample size is small and can reduce the probability of identifying other significant associations or fully uncover the influence of maternal obesity on the bioactive diversity present in samples. Although small cohorts are common in colostrum-related studies due to the ethical and logistical challenges of sample collection, we acknowledge that the implementation of larger sample sizes would improve the creation of integrative datasets, thereby enhancing statistical power and result relevance. Furthermore, while longitudinal data can robust our observations, one of the main purposes of analysing colostrum is to describe the initial biological message that lactating mother has formulated for the neonate. Working with low biomass samples, such as colostrum, presents significant challenges to identify truly relevant signals that may be hidden by contamination noise. On this regard, experimental considerations must be taken for appropriately address the presence of microorganisms in colostrum, including processing in aseptic conditions and in a biological safety cabinet to minimize the number of potential contaminants, and the inclusion of multiple controls to recognize contamination. Additionally, it is important to consider the limitations associated with 16S rDNA assessment. Potential biases introduced by PCR amplification and insufficient sensitivity for detecting deeper taxonomic profiles may result in an incomplete representation of all bacteria in the dataset and significant co-occurrence interactions between species and bioactives may not have been captured. Moreover, breastmilk metabolomics is a developing field with ongoing changes and improvements in analytical technology. The lack of a comprehensive human breastmilk metabolite database or repository hinders the identification and discovery of new metabolites and their origin. Incorporating advanced techniques, such as metagenomic analysis, can offer deeper insights into the functional implications of microbial-host associations in colostrum and their modulation by obesity. Additionally, expanding the scope of immunological assessments to include additional cytokines, such as Th1 and Th2 profiles, as well as immunoglobulins, may substantially enrich our comprehension of colostrum’s immunomodulatory properties and elucidate their functional significance within colostrum. Finally, while co-occurrence networks offer insights into microbial community structure and their interactions with bioactive molecules that traditional methods might overlook, they do not establish directional interactions or causal mechanisms. Co-occurrence analysis relies on correlation coefficients to quantify associations, which may miss nonlinear relationships or interactions mediated by unmeasured variables.

## Conclusions

Despite colostrum is recognized as a dynamic biological system^10^, research into how maternal obesity influences its bioactive profile remains limited. Using co-occurrence networks, we integrated metataxonomic data, metabolomics, and cytokines profile to provide novel insights and a proof of concept for colostrum as an interconnected system and demonstrate the influence of maternal obesity on bioactive interactions within this microenvironment. Our analysis identified a negative co-occurrence between the lipid mediator 15-HEDE and IL-10, while also showing a positive association with microbiota members such as *Beijerinckiaceae* and *Burkholderiaceae* – findings that have not been previously reported in colostrum from mothers with obesity. Additionally, we identified that *Aeromonadaceae*, *Xanthomonadaceae*, and *Staphylococcaceae* were negatively correlated with proinflammatory cytokines (TNF-α, IL-6, and IL-12p70), suggesting a potential regulatory effect on inflammatory responses in colostrum associated with maternal obesity. Our findings reveal that maternal obesity influences alterations in ecosystem structure. However, further research is needed to elucidate the neonatal health implications of the disrupted interactions between bacterial community, metabolites and local inflammatory markers observed in colostrum from women with obesity.

## Author contributions

JSGV: Data curation, Formal analysis, Investigation, Methodology, Writing – original draft, Writing – review & editing. KCC: Data curation, Formal analysis, Investigation, Supervision, Writing – review & editing. SSS: Investigation, Methodology, Writing – review & editing. MRAG: Investigation, Methodology. CNLV: Investigation, Methodology. RACC: Formal analysis, Methodology, Writing – review & editing. AMU: Supervision, Formal analysis, Methodology, Writing – review & editing. VJLD: Investigation, Methodology, Supervision, Writing – review & editing. MB: Conceptualization, Investigation, Methodology, Supervision, Funding acquisition, Writing – original draft, Writing – review & editing. CLC: Conceptualization, Data curation, Formal analysis, Funding acquisition, Project administration, Supervision, Writing – original draft, Writing – review & editing.

## Data availability

All fastq-format reads analyzed have been deposited and are publicly available at the NCBI Sequence Read Archive (SRA). Metabolite raw datasets are available at the GNPS/MassIVE public repository and are available as of the date of publication. Accession numbers are listed in the Supplementary Information (Table S7). This paper does not report original code.

## Acknowledgements

This work was financially supported by Centro de Biotecnología FEMSA from Tecnológico de Monterrey, The Institute for Obesity Research, from Tecnológico de Monterrey. We thank the National Council for Humanities Science and Technology (CONAHCYT) for providing scholarship to JSGV; Frontera de la Ciencia grant No. CF/2023/G/990 to MB, BIOCODEX Microbiota Foundation grant to CLC and grant No. 314964 (Program F0001-2020-02) that supported the functioning of the LC-MS equipment. Finally, we would like to thank Diana Priscila Bonilla Ruelas and Alan G. Hernández Melgar for their technical support.

## Footnotes

Electronic supplementary information (ESI) available.

## References

1 L. F. Stinson, A. S. M. Sindi, A. S. Cheema, C. T. Lai, B. S. Mühlhäusler, M. E. Wlodek, M. S. Payne and D. T. Geddes, The human milk microbiome: who, what, when, where, why, and how?, Nutr. Rev., 2021, 79, 529–543.

2 G. A. G. Lokossou, L. Kouakanou, A. Schumacher and A. C. Zenclussen, Human breast milk: from food to active immune response with disease protection in infants and mothers., Front. Immunol., 2022, 13, 849012.

3 S. Perrella, Z. Gridneva, C. T. Lai, L. Stinson, A. George, S. Bilston-John and D. Geddes, Human milk composition promotes optimal infant growth, development and health., Semin. Perinatol., 2021, 45, 151380.

4 S. Moossavi, S. Sepehri, B. Robertson, L. Bode, S. Goruk, C. J. Field, L. M. Lix, R. J. de Souza, A. B. Becker, P. J. Mandhane, S. E. Turvey, P. Subbarao, T. J. Moraes, D. L. Lefebvre, M. R. Sears, E. Khafipour and M. B. Azad, Composition and Variation of the Human Milk Microbiota Are Influenced by Maternal and Early-Life Factors., Cell Host Microbe, 2019, 25, 324–335.e4.

5 E. Isganaitis, S. Venditti, T. J. Matthews, C. Lerin, E. W. Demerath and D. A. Fields, Maternal obesity and the human milk metabolome: associations with infant body composition and postnatal weight gain., Am. J. Clin. Nutr., 2019, 110, 111–120.

6 M. C. Neville, E. W. Demerath, J. Hahn-Holbrook, R. C. Hovey, J. Martin-Carli, M. A. McGuire, E. R. Newton, K. M. Rasmussen, M. C. Rudolph and D. J. Raiten, Parental factors that impact the ecology of human mammary development, milk secretion, and milk composition-a report from “Breastmilk Ecology: Genesis of Infant Nutrition (BEGIN)” Working Group 1., Am. J. Clin. Nutr., 2023, 117 **Suppl 1**, S11– S27.

7 Y. Wan, J. Jiang, M. Lu, W. Tong, R. Zhou, J. Li, J. Yuan, F. Wang and D. Li, Human milk microbiota development during lactation and its relation to maternal geographic location and gestational hypertensive status., Gut Microbes, 2020, 11, 1438–1449.

8 O. Koren, L. Konnikova, P. Brodin, I. U. Mysorekar and M. C. Collado, The maternal gut microbiome in pregnancy: implications for the developing immune system., Nat. Rev. Gastroenterol. Hepatol., 2024, 21, 35–45.

9 P. Christian, E. R. Smith, S. E. Lee, A. J. Vargas, A. A. Bremer and D. J. Raiten, The need to study human milk as a biological system., Am. J. Clin. Nutr., 2021, 113, 1063–1072.

10 S. M. Donovan, N. Aghaeepour, A. Andres, M. B. Azad, M. Becker, S. E. Carlson, K. M. Järvinen, W. Lin, B. Lönnerdal, C. M. Slupsky, A. L. Steiber and D. J. Raiten, Evidence for human milk as a biological system and recommendations for study design-a report from “Breastmilk Ecology: Genesis of Infant Nutrition (BEGIN)” Working Group 4., Am. J. Clin. Nutr., 2023, 117 **Suppl 1**, S61–S86.

11 R. Piñeiro-Salvador, E. Vazquez-Garza, J. A. Cruz-Cardenas, C. Licona-Cassani, G. García-Rivas, J. Moreno-Vásquez, M. R. Alcorta-García, V. J. Lara-Diaz and M. E. G. Brunck, A cross-sectional study evidences regulations of leukocytes in the colostrum of mothers with obesity., BMC Med., 2022, 20, 388.

12 K. Daiy, V. Harries, K. Nyhan and U. M. Marcinkowska, Maternal weight status and the composition of the human milk microbiome: A scoping review., PLoS ONE, 2022, 17, e0274950.

13 J. S. Gámez-Valdez, J. F. García-Mazcorro, A. H. Montoya-Rincón, D. L. Rodríguez-Reyes, G. Jiménez-Blanco, M. T. A. Rodríguez, R. P.-C. de Vaca, M. R. Alcorta-García, M. Brunck, V. J. Lara-Díaz and C. Licona-Cassani, Differential analysis of the bacterial community in colostrum samples from women with gestational diabetes mellitus and obesity., Sci. Rep., 2021, 11, 24373.

14 J.-W. Ruan, Y.-C. Liao, P.-C. Chen, Y.-J. Chen, Y.-H. Tsai, P.-J. Tsai, Y.-J. Yang, C.-C. Shieh, Y.-C. Lin and C.-Y. Chi, The composition of the maternal breastmilk microbiota influences the microbiota network structure during early infancy., J. Microbiol. Immunol. Infect., DOI:10.1016/j.jmii.2023.07.005.

15 S. Moossavi, F. Atakora, K. Miliku, S. Sepehri, B. Robertson, Q. L. Duan, A. B. Becker, P. J. Mandhane, S. E. Turvey, T. J. Moraes, D. L. Lefebvre, M. R. Sears, P. Subbarao, C. J. Field, L. Bode, E. Khafipour and M. B. Azad, Integrated analysis of human milk microbiota with oligosaccharides and fatty acids in the CHILD cohort., Front. Nutr., 2019, 6, 58.

16 R. M. Pace, J. E. Williams, B. Robertson, K. A. Lackey, C. L. Meehan, W. J. Price, J. A. Foster, D. W. Sellen, E. W. Kamau-Mbuthia, E. W. Kamundia, S. Mbugua, S. E. Moore, A. M. Prentice, D. G. Kita, L. J. Kvist, G. E. Otoo, L. Ruiz, J. M. Rodríguez, R. G. Pareja, M. A. McGuire, L. Bode and M. K. McGuire, Variation in human milk composition is related to differences in milk and infant fecal microbial communities., Microorganisms, DOI:10.3390/microorganisms9061153.

17 J. E. Williams, W. J. Price, B. Shafii, K. M. Yahvah, L. Bode, M. A. McGuire and M. K. McGuire, Relationships among microbial communities, maternal cells, oligosaccharides, and macronutrients in human milk., J. Hum. Lact., 2017, 33, 540– 551.

18 J. A. Siles, M. García-Sánchez and M. Gómez-Brandón, Studying Microbial Communities through Co-Occurrence Network Analyses during Processes of Waste Treatment and in Organically Amended Soils: A Review., Microorganisms, DOI:10.3390/microorganisms9061165.

19 A. Barberán, S. T. Bates, E. O. Casamayor and N. Fierer, Using network analysis to explore co-occurrence patterns in soil microbial communities., ISME J., 2012, 6, 343– 351.

20 World Health Organization, Obesity and overweight, https://www.who.int/news-room/fact-sheets/detail/obesity-and-overweight, (accessed May 2, 2024).

21 E. Bolyen, J. R. Rideout, M. R. Dillon, N. A. Bokulich, C. C. Abnet, G. A. Al-Ghalith, H. Alexander, E. J. Alm, M. Arumugam, F. Asnicar, Y. Bai, J. E. Bisanz, K. Bittinger, A. Brejnrod, C. J. Brislawn, C. T. Brown, B. J. Callahan, A. M. Caraballo-Rodríguez, J. Chase, E. K. Cope, R. Da Silva, C. Diener, P. C. Dorrestein, G. M. Douglas, D. M. Durall, C. Duvallet, C. F. Edwardson, M. Ernst, M. Estaki, J. Fouquier, J. M. Gauglitz, S. M. Gibbons, D. L. Gibson, A. Gonzalez, K. Gorlick, J. Guo, B. Hillmann, S. Holmes, H. Holste, C. Huttenhower, G. A. Huttley, S. Janssen, A. K. Jarmusch, L. Jiang, B. D. Kaehler, K. B. Kang, C. R. Keefe, P. Keim, S. T. Kelley, D. Knights, I. Koester, T. Kosciolek, J. Kreps, M. G. I. Langille, J. Lee, R. Ley, Y.-X. Liu, E. Loftfield, C. Lozupone, M. Maher, C. Marotz, B. D. Martin, D. McDonald, L. J. McIver, A. V. Melnik, J. L. Metcalf, S. C. Morgan, J. T. Morton, A. T. Naimey, J. A. Navas-Molina, L. F. Nothias, S. B. Orchanian, T. Pearson, S. L. Peoples, D. Petras, M. L. Preuss, E. Pruesse, L. B. Rasmussen, A. Rivers, M. S. Robeson, P. Rosenthal, N. Segata, M. Shaffer, A. Shiffer, R. Sinha, S. J. Song, J. R. Spear, A. D. Swafford, L. R. Thompson, P. J. Torres, P. Trinh, A. Tripathi, P. J. Turnbaugh, S. Ul-Hasan, J. J. J. van der Hooft, F. Vargas, Y. Vázquez-Baeza, E. Vogtmann and J. G. Caporaso, Reproducible, interactive, scalable and extensible microbiome data science using QIIME 2., Nat. Biotechnol., 2019, 37, 852–857.

22 B. J. Callahan, P. J. McMurdie, M. J. Rosen, A. W. Han, A. J. A. Johnson and S. P. Holmes, DADA2: High-resolution sample inference from Illumina amplicon data., Nat. Methods, 2016, 13, 581–583.

23 E. Pruesse, C. Quast, K. Knittel, B. M. Fuchs, W. Ludwig, J. Peplies and F. O. Glöckner, SILVA: a comprehensive online resource for quality checked and aligned ribosomal RNA sequence data compatible with ARB., Nucleic Acids Res., 2007, 35, 7188–7196.

24 N. A. Bokulich, B. D. Kaehler, J. R. Rideout, M. Dillon, E. Bolyen, R. Knight, G. A. Huttley and J. Gregory Caporaso, Optimizing taxonomic classification of marker-gene amplicon sequences with QIIME 2’s q2-feature-classifier plugin., Microbiome, 2018, 6, 90.

25 M. C. de Goffau, S. Lager, S. J. Salter, J. Wagner, A. Kronbichler, D. S. Charnock-Jones, S. J. Peacock, G. C. S. Smith and J. Parkhill, Recognizing the reagent microbiome., Nat. Microbiol., 2018, 3, 851–853.

26 Q. Cao, X. Sun, K. Rajesh, N. Chalasani, K. Gelow, B. Katz, V. H. Shah, A. J. Sanyal and E. Smirnova, Effects of rare microbiome taxa filtering on statistical analysis., Front. Microbiol., 2020, 11, 607325.

27 J. Oksanen, F. G. Blanchet, R. Kindt, P. Legendre, P. Minchin, R. O’Hara, G. Simpson, P. Solymos, M. Stevenes and H. Wagner, Vegan: Community Ecology Package, R package, 2022.

28 P. J. McMurdie and S. Holmes, phyloseq: an R package for reproducible interactive analysis and graphics of microbiome census data., PLoS ONE, 2013, 8, e61217.

29 Y. Vázquez-Baeza, M. Pirrung, A. Gonzalez and R. Knight, EMPeror: a tool for visualizing high-throughput microbial community data., Gigascience, 2013, 2, 16.

30 M. I. Love, W. Huber and S. Anders, Moderated estimation of fold change and dispersion for RNA-seq data with DESeq2., Genome Biol., 2014, 15, 550.

31 H. Lin and S. D. Peddada, Analysis of compositions of microbiomes with bias correction., Nat. Commun., 2020, 11, 3514.

32 V. M. Flores-Núñez, D. A. Camarena-Pozos, J. D. Chávez-González, R. Alcalde-Vázquez, M. N. Vázquez-Sánchez, A. G. Hernández-Melgar, J. Xool-Tamayo, A. Moreno-Ulloa and L. P. P. Martínez, Synthetic Communities Increase Microbial Diversity and Productivity of *Agave tequilana* Plants in the Field, Phytobiomes Journal, 2023, 7, 435–448.

33 E. L. Schymanski, J. Jeon, R. Gulde, K. Fenner, M. Ruff, H. P. Singer and J. Hollender, Identifying small molecules via high resolution mass spectrometry: communicating confidence., Environ. Sci. Technol., 2014, 48, 2097–2098.

34 J. D. Holman, D. L. Tabb and P. Mallick, Employing proteowizard to convert raw mass spectrometry data., Curr. Protoc. Bioinformatics, 2014, **46**, 13.24.1–9.

35 T. Pluskal, S. Castillo, A. Villar-Briones and M. Oresic, MZmine 2: modular framework for processing, visualizing, and analyzing mass spectrometry-based molecular profile data., BMC Bioinformatics, 2010, 11, 395.

36 M. Wang, J. J. Carver, V. V. Phelan, L. M. Sanchez, N. Garg, Y. Peng, D. D. Nguyen, J. Watrous, C. A. Kapono, T. Luzzatto-Knaan, C. Porto, A. Bouslimani, A. V. Melnik, M. J. Meehan, W.-T. Liu, M. Crüsemann, P. D. Boudreau, E. Esquenazi, M. Sandoval-Calderón, R. D. Kersten, L. A. Pace, R. A. Quinn, K. R. Duncan, C.-C. Hsu, D. J. Floros, R. G. Gavilan, K. Kleigrewe, T. Northen, R. J. Dutton, D. Parrot, E. E. Carlson, B. Aigle, C. F. Michelsen, L. Jelsbak, C. Sohlenkamp, P. Pevzner, A. Edlund, J. McLean, J. Piel, B. T. Murphy, L. Gerwick, C.-C. Liaw, Y.-L. Yang, H.-U. Humpf, M. Maansson, R. A. Keyzers, A. C. Sims, A. R. Johnson, A. M. Sidebottom, B. E. Sedio, A. Klitgaard, C. B. Larson, C. A. B. P, D. Torres-Mendoza, D. J. Gonzalez, D. B. Silva, L. M. Marques, D. P. Demarque, E. Pociute, E. C. O’Neill, E. Briand, E. J. N. Helfrich, E. A. Granatosky, E. Glukhov, F. Ryffel, H. Houson, H. Mohimani, J. J. Kharbush, Y. Zeng, J. A. Vorholt, K. L. Kurita, P. Charusanti, K. L. McPhail, K. F. Nielsen, L. Vuong, M. Elfeki, M. F. Traxler, N. Engene, N. Koyama, O. B. Vining, R. Baric, R. R. Silva, S. J. Mascuch, S. Tomasi, S. Jenkins, V. Macherla, T. Hoffman, V. Agarwal, P. G. Williams, J. Dai, R. Neupane, J. Gurr, A. M. C. Rodríguez, A. Lamsa, C. Zhang, K. Dorrestein, B. M. Duggan, J. Almaliti and N. Bandeira, Sharing and community curation of mass spectrometry data with Global Natural Products Social Molecular Networking., Nat. Biotechnol., 2016, 34, 828–837.

37 A. T. Aron, E. C. Gentry, K. L. McPhail, L.-F. Nothias, M. Nothias-Esposito, A. Bouslimani, D. Petras, J. M. Gauglitz, N. Sikora, F. Vargas, J. J. J. van der Hooft, M. Ernst, K. B. Kang, C. M. Aceves, A. M. Caraballo-Rodríguez, I. Koester, K. C. Weldon, S. Bertrand, C. Roullier, K. Sun, R. M. Tehan, C. A. Boya P, M. H. Christian, M. Gutiérrez, A. M. Ulloa, J. A. Tejeda Mora, R. Mojica-Flores, J. Lakey-Beitia, V. Vásquez-Chaves, Y. Zhang, A. I. Calderón, N. Tayler, R. A. Keyzers, F. Tugizimana, N. Ndlovu, A. A. Aksenov, A. K. Jarmusch, R. Schmid, A. W. Truman, N. Bandeira, M. Wang and P. C. Dorrestein, Reproducible molecular networking of untargeted mass spectrometry data using GNPS., Nat. Protoc., 2020, 15, 1954–1991.

38 L.-F. Nothias, D. Petras, R. Schmid, K. Dührkop, J. Rainer, A. Sarvepalli, I. Protsyuk, M. Ernst, H. Tsugawa, M. Fleischauer, F. Aicheler, A. A. Aksenov, O. Alka, P.-M. Allard, A. Barsch, X. Cachet, A. M. Caraballo-Rodriguez, R. R. Da Silva, T. Dang, N. Garg, J. M. Gauglitz, A. Gurevich, G. Isaac, A. K. Jarmusch, Z. Kameník, K. B. Kang, N. Kessler, I. Koester, A. Korf, A. Le Gouellec, M. Ludwig, C. Martin H, L.-I. McCall, J. McSayles, S. W. Meyer, H. Mohimani, M. Morsy, O. Moyne, S. Neumann, H. Neuweger, N. H. Nguyen, M. Nothias-Esposito, J. Paolini, V. V. Phelan, T. Pluskal, R. A. Quinn, S. Rogers, B. Shrestha, A. Tripathi, J. J. J. van der Hooft, F. Vargas, K. C. Weldon, M. Witting, H. Yang, Z. Zhang, F. Zubeil, O. Kohlbacher, S. Böcker, T. Alexandrov, N. Bandeira, M. Wang and P. C. Dorrestein, Feature-based molecular networking in the GNPS analysis environment., Nat. Methods, 2020, 17, 905–908.

39 L. Cao, M. Guler, A. Tagirdzhanov, Y.-Y. Lee, A. Gurevich and H. Mohimani, MolDiscovery: learning mass spectrometry fragmentation of small molecules., Nat. Commun., 2021, 12, 3718.

40 H. Mohimani, A. Gurevich, A. Shlemov, A. Mikheenko, A. Korobeynikov, L. Cao, E. Shcherbin, L.-F. Nothias, P. C. Dorrestein and P. A. Pevzner, Dereplication of microbial metabolites through database search of mass spectra., Nat. Commun., 2018, 9, 4035.

41 K. Dührkop, M. Fleischauer, M. Ludwig, A. A. Aksenov, A. V. Melnik, M. Meusel, P. C. Dorrestein, J. Rousu and S. Böcker, SIRIUS 4: a rapid tool for turning tandem mass spectra into metabolite structure information., Nat. Methods, 2019, 16, 299–302.

42 M. Ludwig, L.-F. Nothias, K. Dührkop, I. Koester, M. Fleischauer, M. A. Hoffmann, D. Petras, F. Vargas, M. Morsy, L. Aluwihare, P. C. Dorrestein and S. Böcker, Database-independent molecular formula annotation using Gibbs sampling through ZODIAC, *Nat*. Mach. Intell., 2020, 2, 629–641.

43 K. Dührkop, H. Shen, M. Meusel, J. Rousu and S. Böcker, Searching molecular structure databases with tandem mass spectra using CSI:FingerID., Proc Natl Acad Sci USA, 2015, 112, 12580–12585.

44 J. Wandy, Y. Zhu, J. J. J. van der Hooft, R. Daly, M. P. Barrett and S. Rogers, Ms2lda.org: web-based topic modelling for substructure discovery in mass spectrometry., Bioinformatics, 2018, 34, 317–318.

45 J. Willforss, A. Chawade and F. Levander, NormalyzerDE: Online Tool for Improved Normalization of Omics Expression Data and High-Sensitivity Differential Expression Analysis., J. Proteome Res., 2019, 18, 732–740.

46 M. E. Ritchie, B. Phipson, D. Wu, Y. Hu, C. W. Law, W. Shi and G. K. Smyth, limma powers differential expression analyses for RNA-sequencing and microarray studies., Nucleic Acids Res., 2015, 43, e47.

47 K. Blighe, S. Rana and M. Lewis, EnhancedVolcano: Publication-ready volcano plots with enhanced colouring andlabeling, Bioconductor, DOI:10.18129/b9.bioc.enhancedvolcano.

48 F. R. Passos Mansoldo, R. Garrett, V. da Silva Cardoso, M. A. Alves and A. B. Vermelho, Metabology: Analysis of metabolomics data using community ecology tools., Anal. Chim. Acta, 2022, 1232, 340469.

49 J. Chong, P. Liu, G. Zhou and J. Xia, Using MicrobiomeAnalyst for comprehensive statistical, functional, and meta-analysis of microbiome data., Nat. Protoc., 2020, 15, 799–821.

50 J. T. Smilowitz, L. H. Allen, D. C. Dallas, J. McManaman, D. J. Raiten, M. Rozga, D. A. Sela, A. Seppo, J. E. Williams, B. E. Young and M. K. McGuire, Ecologies, synergies, and biological systems shaping human milk composition-a report from “Breastmilk Ecology: Genesis of Infant Nutrition (BEGIN)” Working Group 2., Am. J. Clin. Nutr., 2023, 117 **Suppl 1**, S28–S42.

51 S. Enstad, S. Cheema, R. Thomas, R. N. Fichorova, C. R. Martin, P. O’Tierney-Ginn, C. L. Wagner and S. Sen, The impact of maternal obesity and breast milk inflammation on developmental programming of infant growth., Eur. J. Clin. Nutr., 2021, 75, 180–188.

52 H. Nuss, A. Altazan, J. Zabaleta, M. Sothern and L. Redman, Maternal pre-pregnancy weight status modifies the influence of PUFAs and inflammatory biomarkers in breastmilk on infant growth., PLoS ONE, 2019, 14, e0217085.

53 C. R. Sims, M. E. Lipsmeyer, D. E. Turner and A. Andres, Human milk composition differs by maternal BMI in the first 9 months postpartum., Am. J. Clin. Nutr., 2020, 112, 548–557.

54 C. Gómez-Gallego, J. M. Morales, D. Monleón, E. du Toit, H. Kumar, K. M. Linderborg, Y. Zhang, B. Yang, E. Isolauri, S. Salminen and M. C. Collado, Human Breast Milk NMR Metabolomic Profile across Specific Geographical Locations and Its Association with the Milk Microbiota., Nutrients, DOI:10.3390/nu10101355.

55 K. E. Lyons, C.-A. O. ’ Shea, G. Grimaud, C. A. Ryan, E. Dempsey, A. L. Kelly, R. P. Ross and C. Stanton, The human milk microbiome aligns with lactation stage and not birth mode., Sci. Rep., 2022, 12, 5598.

56 G. E. Leghi, P. F. Middleton and B. S. Muhlhausler, A methodological approach to identify the most reliable human milk collection method for compositional analysis: a systematic review protocol., Syst. Rev., 2018, 7, 122.

57 M. S. Matchado, M. Lauber, S. Reitmeier, T. Kacprowski, J. Baumbach, D. Haller and M. List, Network analysis methods for studying microbial communities: A mini review., Comput. Struct. Biotechnol. J., 2021, 19, 2687–2698.

58 M. Layeghifard, D. M. Hwang and D. S. Guttman, Disentangling interactions in the microbiome: A network perspective., Trends Microbiol., 2017, 25, 217–228.

59 M. Loftus, S. A.-D. Hassouneh and S. Yooseph, Bacterial associations in the healthy human gut microbiome across populations., Sci. Rep., 2021, 11, 2828.

60 C. J. Robinson, B. J. M. Bohannan and V. B. Young, From structure to function: the ecology of host-associated microbial communities., Microbiol. Mol. Biol. Rev., 2010, 74, 453–476.

61 D. J. Lemas, B. E. Young, P. R. Baker, A. C. Tomczik, T. K. Soderborg, T. L. Hernandez, B. A. de la Houssaye, C. E. Robertson, M. C. Rudolph, D. Ir, Z. W. Patinkin, N. F. Krebs, S. A. Santorico, T. Weir, L. A. Barbour, D. N. Frank and J. E. Friedman, Alterations in human milk leptin and insulin are associated with early changes in the infant intestinal microbiome., Am. J. Clin. Nutr., 2016, 103, 1291– 1300.

62 S. Beretta, M. Apparicio, G. H. Toniollo and M. V. Cardozo, The importance of the intestinal microbiota in humans and dogs in the neonatal period., Anim. Reprod., 2023, 20, e20230082.

63 C. D. Moon, W. Young, P. H. Maclean, A. L. Cookson and E. N. Bermingham, Metagenomic insights into the roles of Proteobacteria in the gastrointestinal microbiomes of healthy dogs and cats., Microbiologyopen, 2018, 7, e00677.

64 U. Pandey and P. Aich, Postnatal intestinal mucosa and gut microbial composition develop hand in hand: A mouse study., Biomed. J., 2023, 46, 100519.

65 K. Hou, Z.-X. Wu, X.-Y. Chen, J.-Q. Wang, D. Zhang, C. Xiao, D. Zhu, J. B. Koya, L. Wei, J. Li and Z.-S. Chen, Microbiota in health and diseases., Signal Transduct. Target. Ther., 2022, 7, 135.

66 J. Mirpuri, M. Raetz, C. R. Sturge, C. L. Wilhelm, A. Benson, R. C. Savani, L. V. Hooper and F. Yarovinsky, Proteobacteria-specific IgA regulates maturation of the intestinal microbiota., Gut Microbes, 2014, 5, 28–39.

67 E. Cortés-Macías, M. Selma-Royo, K. Rio-Aige, C. Bäuerl, M. J. Rodríguez-Lagunas, C. Martínez-Costa, F. J. Pérez-Cano and M. C. Collado, Distinct breast milk microbiota, cytokine, and adipokine profiles are associated with infant growth at 12 months: an in vitro host-microbe interaction mechanistic approach., Food Funct., 2023, 14, 148–159.

68 A. Köhler, S. Delbauve, J. Smout, D. Torres and V. Flamand, Very early-life exposure to microbiota-induced TNF drives the maturation of neonatal pre-cDC1., Gut, 2021, 70, 511–521.

69 E. Amabebe and D. O. Anumba, Diabetogenically beneficial gut microbiota alterations in third trimester of pregnancy, Reproduction and Fertility, 2021, 2, R1– R12.

70 M. C. Collado, K. Laitinen, S. Salminen and E. Isolauri, Maternal weight and excessive weight gain during pregnancy modify the immunomodulatory potential of breast milk., Pediatr. Res., 2012, 72, 77–85.

71 F. Bardanzellu, M. Puddu, D. G. Peroni and V. Fanos, The human breast milk metabolome in overweight and obese mothers., Front. Immunol., 2020, 11, 1533.

72 J. L. Saben, C. R. Sims, B. D. Piccolo and A. Andres, Maternal adiposity alters the human milk metabolome: associations between nonglucose monosaccharides and infant adiposity., Am. J. Clin. Nutr., 2020, 112, 1228–1239.

73 H. H. Chen, Y. J. Tseng, S. Y. Wang, Y. S. Tsai, C. S. Chang, T. C. Kuo, W. J. Yao, C. C. Shieh, C. H. Wu and P. H. Kuo, The metabolome profiling and pathway analysis in metabolic healthy and abnormal obesity., Int J Obes (Lond), 2015, 39, 1241–1248.

74 T. Purdom, L. Kravitz, K. Dokladny and C. Mermier, Understanding the factors that effect maximal fat oxidation., J. Int. Soc. Sports Nutr., 2018, 15, 3.

75 M. Zhang, L. Simon Sarkadi, M. Üveges, J. Tormási, E. Benes, R. A. Vass and S. G. Vari, Gas chromatographic determination of fatty acid composition in breast milk of mothers with different health conditions, Acta Alimentaria, 2022, 51, 625–635.

76 R. E. Walker, Oxylipins as potential regulators of inflammatory conditions of human lactation., Metabolites, DOI:10.3390/metabo12100994.

77 G. A. Weiss, H. Troxler, G. Klinke, D. Rogler, C. Braegger and M. Hersberger, High levels of anti-inflammatory and pro-resolving lipid mediators lipoxins and resolvins and declining docosahexaenoic acid levels in human milk during the first month of lactation., Lipids Health Dis., 2013, 12, 89.

78 R. Ottria, M. D. Porta, O. Xynomilakis, S. Casati, R. Cazzola and P. Ciuffreda, Lipids and lipid signaling molecules in Human milk and infant formula, a chemical characterization of relevant biochemical components., J. Nutr. Biochem., 2024, 109580.

79 B. Koletzko and M. Rodriguez-Palmero, Polyunsaturated fatty acids in human milk and their role in early infant development., J. Mammary Gland Biol. Neoplasia, 1999, 4, 269–284.

80 A. D. George, S. Burugupalli, S. Paul, T. Mansell, D. Burgner and P. J. Meikle, The Role of Human Milk Lipids and Lipid Metabolites in Protecting the Infant against Non-Communicable Disease., Int. J. Mol. Sci., DOI:10.3390/ijms23147490.

81 J. Kolmert, S. Piñeiro-Hermida, M. Hamberg, J. A. Gregory, I. P. López, A. Fauland, C. E. Wheelock, S.-E. Dahlén, J. G. Pichel and M. Adner, Prominent release of lipoxygenase generated mediators in a murine house dust mite-induced asthma model., Prostaglandins Other Lipid Mediat., 2018, 137, 20–29.

82 Y.-S. Huang, W.-C. Huang, C.-W. Li and L.-T. Chuang, Eicosadienoic acid differentially modulates production of pro-inflammatory modulators in murine macrophages., Mol. Cell. Biochem., 2011, 358, 85–94.

83 F. Haviv, J. D. Ratajczyk, R. W. DeNet, Y. C. Martin, R. D. Dyer and G. W. Carter, Structural requirements for the inhibition of 5-lipoxygenase by 15-hydroxyeicosa-5,8,11,13-tetraenoic acid analogues., J. Med. Chem., 1987, 30, 254–263.

84 K. Miyata, D. Horikami, Y. Tachibana, T. Yamamoto, T. Nakamura, K. Kobayashi and T. Murata, 15-hydroxy eicosadienoic acid is an exacerbating factor for nasal congestion in mice., FASEB J., 2022, 36, e22085.

85 T. Takeichi, F. Kinoshita, H. Tanaka, S. Fujita, Y. Kobayashi, M. Nakatochi, K. Sugiura and M. Akiyama, The lipoxygenaseChepoxilin pathway is activated in cutaneous plaque lesions of psoriasis, J. Cutan. Immunol. Allergy, 2019, 2, 15–24.

86 K. Esposito, A. Pontillo, F. Giugliano, G. Giugliano, R. Marfella, G. Nicoletti and D. Giugliano, Association of low interleukin-10 levels with the metabolic syndrome in obese women., J. Clin. Endocrinol. Metab., 2003, 88, 1055–1058.

87 N. Subramanian, B. Tavira, K. Hofwimmer, B. Gutsmann, L. Massier, J. Abildgaard, A. Juul, M. Rydén, P. Arner and J. Laurencikiene, Sex-specific regulation of IL-10 production in human adipose tissue in obesity., Front Endocrinol (Lausanne), 2022, 13, 996954.

88 V. J. Taylor, Lactation from the inside out: Maternal homeorhetic gastrointestinal adaptations regulating energy and nutrient flow into milk production., Mol. Cell. Endocrinol., 2023, 559, 111797.

89 F. Kalaycı-Yüksek, D. Gümüş, V. Güler, A. Uyanık-Öcal and M. Anğ-Küçüker, Progesterone and Estradiol alter the growth, virulence and antibiotic susceptibilities of Staphylococcus aureus., New Microbiol., 2023, 46, 43–51.

90 M. Kisiela, A. Skarka, B. Ebert and E. Maser, Hydroxysteroid dehydrogenases (HSDs) in bacteria: a bioinformatic perspective., J. Steroid Biochem. Mol. Biol., 2012, 129, 31–46.

91 M. Nuriel-Ohayon, H. Neuman, O. Ziv, A. Belogolovski, Y. Barsheshet, N. Bloch, A. Uzan, R. Lahav, A. Peretz, S. Frishman, M. Hod, E. Hadar, Y. Louzoun, O. Avni and O. Koren, Progesterone Increases Bifidobacterium Relative Abundance during Late Pregnancy., Cell Rep., 2019, 27, 730–736.e3.

92 I. Bartha, I. Joumady, M. Cuerva and J. L. Bartha, The effect of maternal obesity and lipid profile on first-trimester serum progesterone levels., American Journal of Obstetrics & Gynecology MFM, 2023, 5, 100959.

93 M. Lu, H. Xiao, K. Li, J. Jiang, K. Wu and D. Li, Concentrations of estrogen and progesterone in breast milk and their relationship with the mother’s diet., Food Funct., 2017, 8, 3306–3310.

94 M. Messripour, A. Forooghi-Abary and F. Dashti, Association between sex hormones in human breast milk and infant growth and development, Arch Iranian Med., 2002, 5, 166-169.

