## Supplementary Figures for "Analysis of human colostrum reveals differential co-occurrence networks of metabolites, microbiota and cytokines in maternal obesity"

**Supplementary Information**

**Ecology-based analysis of human colostrum unearth differential co-occurrence networks of metabolites, microbiota and cytokines in maternal obesity: A proof of concept**

July S. Gámez-Valdez^a,b^, Karina Corona-Cervantes^b^, Erick S. Sanchez-Salguero^c^, Mario R. Alcorta-García^d,e^, Claudia N. Lopez-Villaseñor^d,f^, Rommel A. Carballo-Castañeda^g^, Aldo Moreno-Ulloa^g^, Victor J. Lara-Diaz^f,h^, Marion E. G. Brunck^a,c^ and Cuauhtémoc Licona-Cassani ^a,b*^

^a^ Centro de Biotecnología FEMSA, Escuela de Ingeniería y Ciencias, Tecnologico de Monterrey, N.L. México.

^b^ Unidad de Biología Integrativa, The Institute for Obesity Research, Tecnologico de Monterrey, N.L. México.

^c^ Unidad de Biología Experimental, The Institute for Obesity Research, Tecnologico de Monterrey, N.L. México.

^d^ Hospital Regional Materno Infantil, Servicios de Salud de Nuevo León, OPD, Ciudad Guadalupe, N.L. México.

^e^ Hospital San José TECSALUD, Monterrey, N.L. México.

^f^ Escuela de Medicina y Ciencias de la Salud, Tecnológico de Monterrey, N.L. México.

^g^ Departamento de Innovación Biomédica, Centro de Investigación Científica y de Educación Superior de Ensenada, Baja California (CICESE), Ensenada, México.

^h^ Faculty of Medicine, The University of New South Wales, Sydney, Australia.

**Table of contents - Figures**

**Figure S1.**.	2

**Figure S2.**	3

**Figure S3.**.	4

**Figure S4.**	5

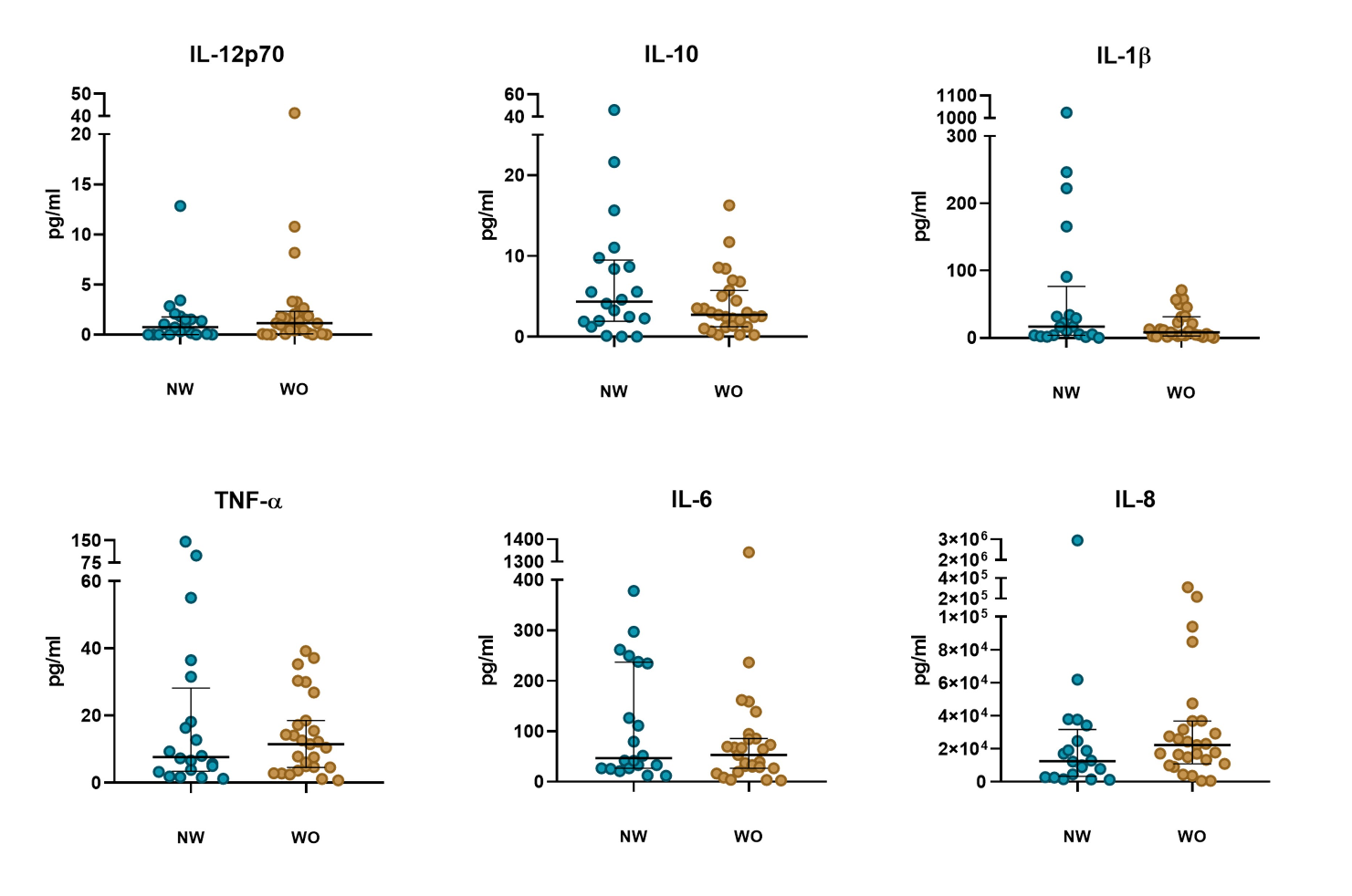

**Figure S1.** Cytokine concentrations in the colostrum from the normal weight (NW; blue) and with obesity (WO; brown) group, related to Table 1. Comparisons performed using Mann-Whitney´s U tests. Median values and interquartile (IQ) range are represented.

**Figure S2.**
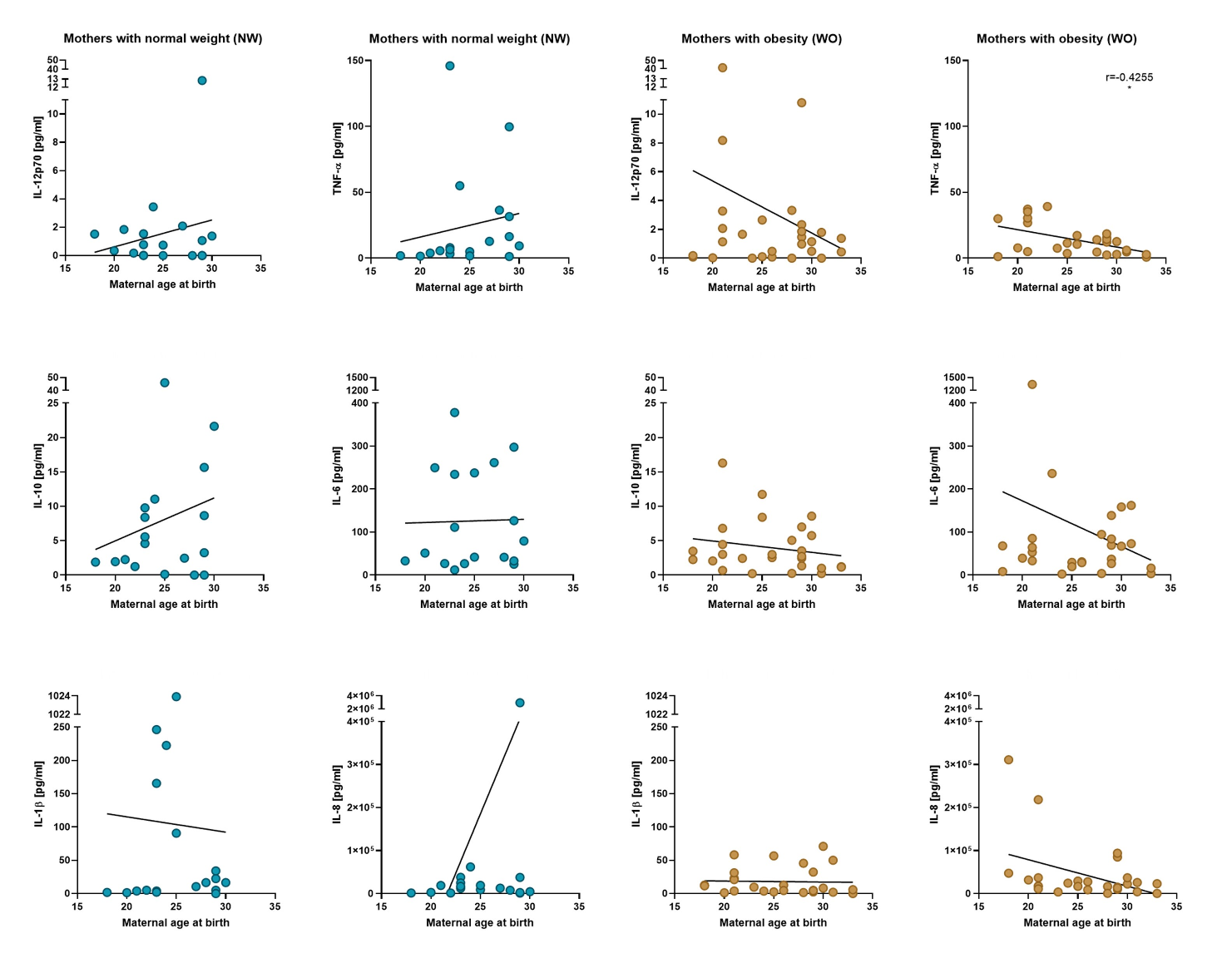
Correlations between maternal age and cytokine concentrations in the colostrum from the normal weight (blue) and with obesity group (brown), related to Table 1. Trends were analyzed with Spearman’s test.

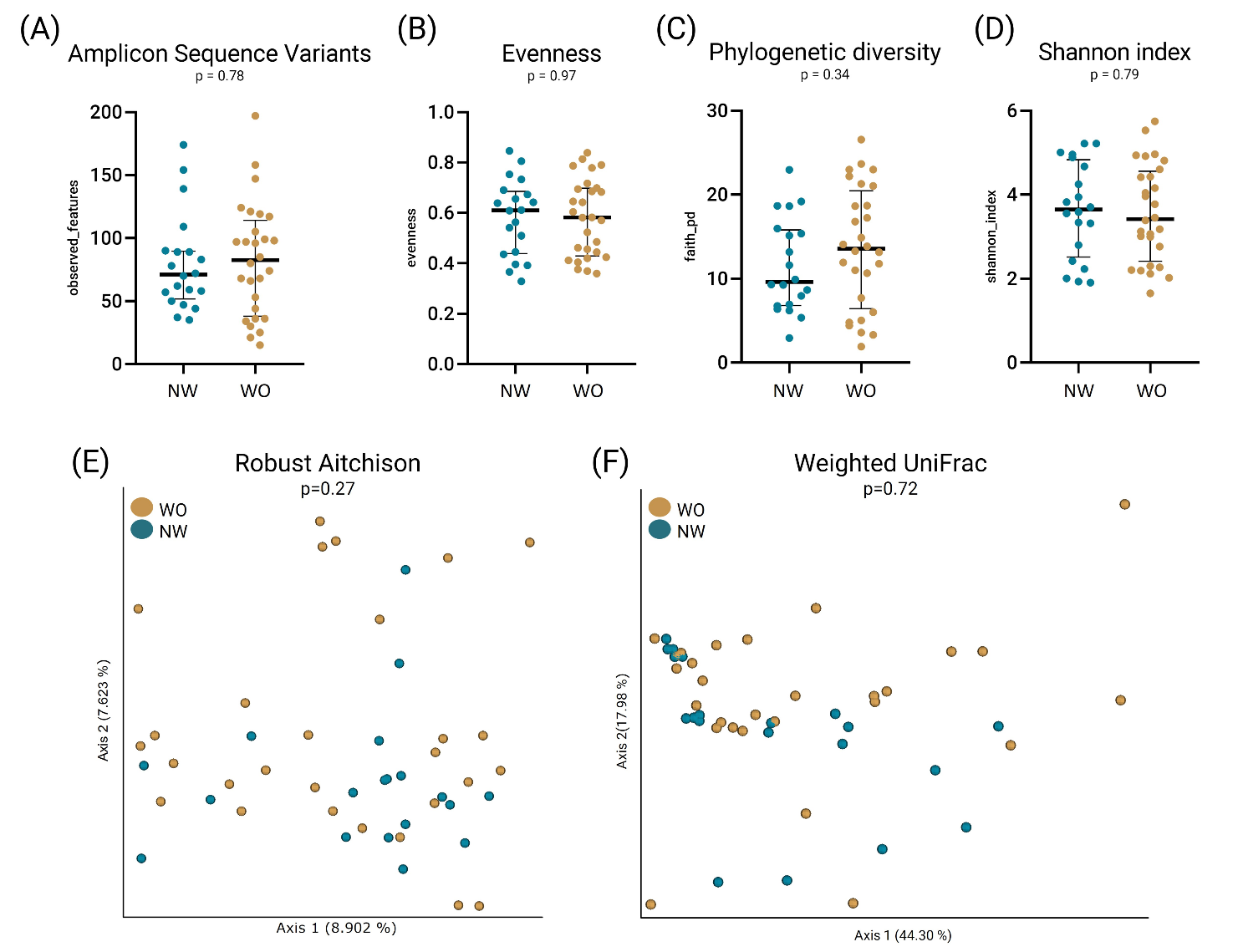

**Figure S3.** Colostrum microbiota diversity. (A-D) Alpha diversity: (A) observed amplicon sequences variants (ASVs); (B) Evenness; (C) Phylogenetic distance; (D) Shannon index; (E-F) Beta diversity: (E) Robust Principal Component Analysis (RPCA) from the Robust Aitchison distance; (F) Principal Coordinates Analysis (PCoA) from the Weighted UniFrac distance. Beta diversity indexes were visualized using EMPEROR. NW, normal weight group; WO, with obesity group.

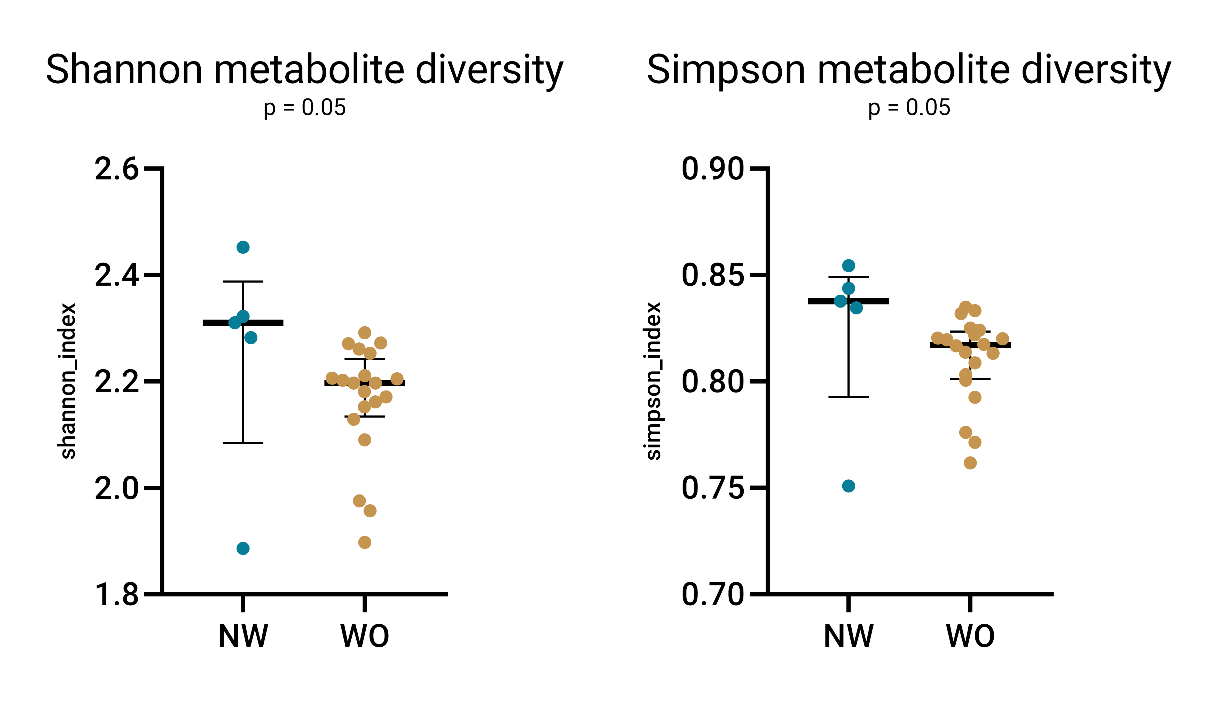

**Figure S4.** Chemical diversity of colostrum among groups, related to Figure 2. (A-B) Boxplots of the chemical diversity of samples using (A) Shannon index and (B) Simpson index. Comparison was evaluated through Wilcoxon rank sum test; NW, normal weight group; WO, with obesity group.
